# How cells determine the number of polarity sites

**DOI:** 10.1101/2020.05.21.109520

**Authors:** Jian-geng Chiou, Kyle D. Moran, Daniel J. Lew

## Abstract

The diversity of cell morphologies arises, in part, through regulation of cell polarity by Rho-family GTPases. A poorly understood but fundamental question concerns the regulatory mechanisms by which different cells can generate different numbers of polarity sites. Theoretical models of polarity circuits develop multiple initial polarity sites, but then those sites engage in competition, leaving a single winner. The timescale of competition slows dramatically as GTPase concentrations at polarity sites approach a “saturation point”, allowing multiple sites to coexist. Here, we show that these principles hold in more complex mechanistic models of the *Saccharomyces cerevisiae* polarity machinery, and confirm model predictions *in vivo*. Further, we elucidate a novel design principle whereby cells can switch from competition to equalization among polarity sites. These findings provide insight into how cells determine the number of polarity sites.

## Introduction

Eukaryotic cells display a very wide diversity of cell morphologies, which are often critical to carry out specialized cell functions. Different morphologies arise through specific arrangements and actions of the cytoskeleton. In turn, the cytoskeleton is regulated by the conserved Rho family of GTPases (Etienne-Manneville and Hall, 2002). These GTPases act as molecular switches, active when bound to GTP and inactive when bound to GDP, that can associate with cell membranes through hydrophobic lipid modifications. Switching from GDP-to GTP-bound forms is catalyzed by guanine nucleotide exchange factors (GEFs), while switching back from GTP-to GDP-bound forms is catalyzed by GTPase activating proteins (GAPs). A subset of these GTPases (Cdc42, Rac, Rop) regulates cell polarity (Park and Bi, 2007; Wu and Lew, 2013). Active polarity GTPases become concentrated at one or more regions of the plasma membrane, where they can bind and recruit effector proteins that regulate the cytoskeleton as well as vesicle traffic. In many cell types (e.g. migrating cells, plant pollen tubes and root hairs, or several budding yeasts), it is crucial to maintain one and only one polarity domain, establishing a single polarity axis (front) that leads to movement or growth in that direction (Chiou et al., 2017; Houk et al., 2012; Wu and Lew, 2013; Yang and Lavagi, 2012). In other cell types (e.g. neurons with many neurite tips, plant cells that form xylem, or filamentous fungal cells with branches), multiple active-GTPase clusters coexist in the same cell (Dotti et al., 1988; Knechtle et al., 2003; Oda and Fukuda, 2012). These differences raise the question of how Rho-GTPase polarity systems in specific cell types can be tuned to yield the desired number of polarized fronts.

Key properties of pattern formation by polarity GTPase systems can be captured by mathematical models of a class we will refer to as Mass Conserved Activator Substrate (MCAS) models (Chiou et al., 2018; Goryachev and Pokhilko, 2008; Halatek et al., 2018; Jilkine and Edelstein-Keshet, 2011; Mori et al., 2008; Otsuji et al., 2007; Otsuji et al., 2010). These models consist of sets of partial differential equations (PDEs) that encode the interconversion of polarity factors between two forms: a membrane-bound (and hence slow-diffusing) “activator” and a cytosolic (and hence rapidly-diffusing) “substrate” (Fig. 1A). One critical feature of these systems is positive feedback, such that membrane regions with higher concentrations of activator can locally recruit and activate more substrate from the cytoplasm. A second critical feature is the difference in diffusivity between the activator and the substrate, allowing a localized accumulation of activator to recruit substrate from a much wider region of cytoplasm. A third critical feature is mass conservation: because the combined amount of activator and substrate is fixed, accumulation of activator at the membrane depletes substrate from the cytoplasm, limiting the size, activator concentration, and potentially the number of permissible activator-enriched regions.

**Figure 1.**
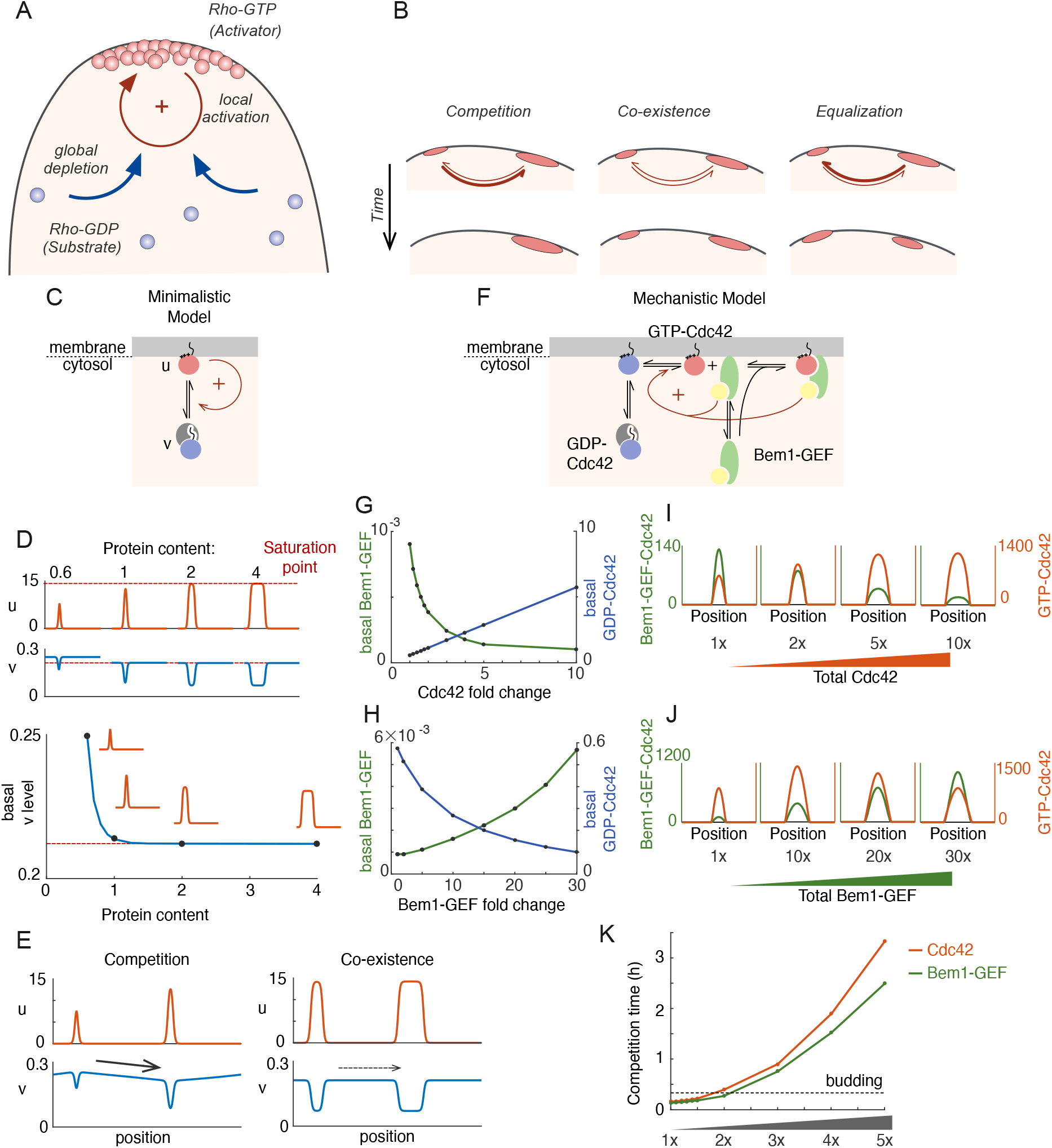
Competition in a mechanistic Rho-GTPase model. A) Rho-GTPase polarity circuits can be modeled as mass-conserved reaction-diffusion systems where membrane-bound Rho-GTP is a slow-diffusing activator and cytosolic Rho-GDP is a fast-diffusing substrate. Such systems polarize Rho-GTPase to spatially confined polarity sites based on positive feedback (+) via local activation that recruits substrate from the cytoplasm, leading to global substrate depletion. B) A starting condition with two unequal patches of activator can evolve in three ways. Competition occurs if the larger patch recruits substrates more than the smaller; coexistence occurs if both patches recruit substrate equally well; and equalization occurs if the smaller patch recruits substrates more than the larger. C) Schematic of the minimalistic MCAS model with one Rho-GTPase converting between activator (u: Rho-GTP) and substrate (v: Rho-GDP) forms. Arrows depict reactions. The positive feedback is highlighted in red (see text for details). D) In the minimalistic model, as total protein content in the system increases, peak u concentration approaches a saturation point, while basal v concentration declines to a limit. 1x protein content equivalent to starting uniform v concentration of 2. E) Snapshots of simulations starting from two single peak steady states placed next to each other in the same domain. With patches far from saturation, the larger patch depletes more substrate than the smaller (Left panel, starting protein contents 0.6x, 1x), leading to a net flux (arrow) that results in competition. With patches close to saturation, substrate depletion is similar for both (Right panel, starting protein contents 2x, 4x), leading to little flux and therefore coexistence. F) Schematic of a mechanistic model of the yeast polarity circuit: arrows depict reactions assumed to occur with mass-action kinetics. The positive feedback is highlighted in red. G,H) Basal concentrations of the cytoplasmic substrates GDP-Cdc42 and Bem1-GEF as a function of total Cdc42 (G) or Bem1-GEF (H) content. I,J) Membrane protein concentration profiles of the activators GTP-Cdc42 and the Bem1-GEF-Cdc42 complex as a function of total Cdc42 (I) or Bem1-GEF (J) content. Note that the concentration of all GTP-Cdc42 in the peak is the sum of unbound (GTP-Cdc42) and bound (Bem1-GEF-Cdc42) protein. K) Time taken to resolve competition as Cdc42 or Bem1-GEF content is increased. Timescale in hours. Dashed line at 20 min indicates time from cell cycle start to budding in *Saccharomyces cerevisiae*. Total protein content in panels G-I: 1x Cdc42 corresponds to 1 μM in the cell; 1x Bem1-GEF corresponds to 0.017 μM in the cell.

MCAS models at the homogeneous steady state can develop inhomogeneous activator distributions either spontaneously through Turing instability, or in response to external cues. Once a local region becomes enriched for activator, it grows (acquires more activator) by recruiting more activator from the cytoplasm, eventually depleting cytoplasmic substrate levels until the system reaches a polarized steady state with a local peak in activator concentration. However, the fate of any given peak depends on the presence of other peaks, which can also deplete substrate from the cytoplasm. Clusters (peaks) that differ in total protein content would also differ in their ability to recruit cytoplasmic substrate (Chiou et al., 2018). A hypothetical case with two initial unequal peaks could evolve in three possible directions (Fig. 1B): 1) Competition: If the peak with greater activator content grows faster than the smaller peak, then as cytoplasmic substrate levels become depleted, the smaller peak would be starved of the fuel it needs to survive, and begin to shrink as loss of activator to the cytoplasm outpaces recruitment of fresh substrate. The shrinking of the smaller peak would restore substrate to the cytoplasm, allowing the larger peak to grow further until there is only one peak at steady state. 2) Co-existence: If the two unequal peaks both grow at the same rate, then their growth would slow in parallel as the substrate is depleted from the cytoplasm. When substrate depletion makes recruitment slow enough to match the rate at which activator is lost from the peaks back to the cytoplasm, the unequal peaks would persist indefinitely. 3) Equalization: If the smaller peak grows more rapidly than the larger peak, then that would continue until the two are equal.

Previous work on minimalistic one-activator, one-substrate MCAS models indicated that the mass conservation feature enforces the competition scenario, with the largest peak becoming the only one (Chiou et al., 2018; Otsuji et al., 2007). However, our recent work has shown that the growth rate of a peak (for a given concentration of available substrate) “saturates” as the peak exceeds a certain activator content. If more than one peak approach this saturation point, then the system switches to a co-existence scenario for biologically relevant timescales. This conclusion is general for minimalistic MCAS models, and the degree of saturation is the dominant factor determining uni- or multi-polar outcomes (Chiou et al., 2018). Equalization does not appear possible in minimalist MCAS models but may be possible in more complicated ones.

One well-studied example of an MCAS system applies to the budding yeast *Saccharomyces cerevisiae*, where the polarity GTPase Cdc42 cycles between a slow-diffusing GTP-bound form and a rapidly diffusing GDP-bound form. GTP-Cdc42 binds effector p21-activated kinases (PAKs), which bind the scaffold protein Bem1, which binds the GEF Cdc24 that activates Cdc42 (Johnson et al., 2011; Kozubowski et al., 2008). This set of interactions allows GTP-Cdc42 to recruit its own GEF, activating neighboring Cdc42 to yield positive feedback. Competition has been observed experimentally in this system, leading to development of a single Cdc42-enriched cortical region that generates a single bud in every cell cycle (Howell et al., 2012; Wu et al., 2015). In addition to positive feedback, the budding yeast Cdc42 system also displays negative feedback loops thought to be mediated by inhibitory GEF phosphorylation (Kuo et al., 2014) and local recruitment of GAPs (Okada et al., 2013). Interestingly, modeling of multi-component systems with negative as well as positive feedback led to competition in some parameter regimes, but equalization in other parameter regimes (Howell et al., 2012; Jacobs et al., 2019).

It seems likely that cells that reliably develop a single polarity site operate with a polarity circuit that always yields competition, whereas cells that develop two or more polarity sites operate with a polarity circuit that yields either coexistence or equalization. There are also systems, like the fission yeast *Schizosaccharomyces pombe*, that switch in a programmed manner between unipolar and bipolar outcomes (Martin and Chang, 2005). It is unclear whether circuits that produce different outcomes differ in their wiring (the reactions undertaken by the polarity factors) or simply in some relevant parameter (e.g. protein amount). Here we identify the features of mechanistic models of the budding yeast Cdc42 system that could lead to co-existence or equalization, and explore the predictions experimentally.

## Results

### A mechanistic model for the yeast Cdc42 system exhibits saturation, and switches from competition to coexistence as protein levels increase

Minimalistic models are intended to capture the essential elements and behaviors of a complex system. Their simplicity can yield conceptual insight, and allows an in-depth analysis of behaviors that would not be feasible for more complex models. However, it can be challenging to translate these concepts into biologically testable hypotheses, as model species and reactions generally represent complex combinations of a cell’s components and biochemical reactions. More complex mechanistic models are harder to analyze mathematically, but they better represent a subset of the real biological species and reactions, so that modeling predictions are testable experimentally. Moreover, the added complexity of mechanistic models can produce unexpected deviations from the behavior exhibited by minimalistic models. As a first step to generate testable predictions, we asked whether a mechanistic model of the budding yeast polarity circuit behaved in a similar way to the minimalistic model in terms of saturation.

In minimalistic models, peaks with greater amout of activator always grow faster than peaks with less activator, but growth rate saturates above a certain activator content (Fig. 1C)(Chiou et al., 2018). The greater the amount of activator in a peak, the more it depletes the cytoplasmic substrate at steady state, until a plateau is reached (Fig. 1D). Thus, when two peaks of unequal activator content are present, the larger peak exerts a greater depletion of substrate, creating a cytoplasmic substrate gradient towards the larger peak that drives competition. However, this gradient becomes negligible between two saturated peaks, resulting in apparent co-existence (Fig. 1E). This general result applies to both 1D and 2D models, although timescales of competition may differ (Chiou et al., 2018).

We first asked whether a more complex mechanistic 2D model of the yeast system behaves in a similar manner to the minimalistic models. This model includes a Bem1-GEF complex as well as Cdc42 (Fig. 1F)(Goryachev and Pokhilko, 2008; Howell et al., 2012): it has two species analogous to “activators” at the membrane (GTP-Cdc42 and the Bem1-GEF-Cdc42 complex) that promote positive feedback, and two species analogous to “substrates” in the cytoplasm (GDP-Cdc42 and Bem1-GEF), as well as other species with different characteristics (GDP-Cdc42 and Bem1-GEF at the membrane). As the total Cdc42 in the system was increased, cytoplasmic Bem1-GEF levels were depleted but Cdc42-GDP accumulated (Fig. 1G). This suggested that the less abundant Bem1-GEF is a limiting substrate, while Cdc42 is present in excess. If instead the total Bem1-GEF was increased, cytoplasmic levels of Bem1-GEF rose, while cytoplasmic Cdc42-GDP became depleted (Fig. 1H). This suggested that the limiting substrate switched from Bem1-GEF to GDP-Cdc42. Although the cytoplasmic level of the limiting substrate decreased as the amount of the other substrate increased, the level of the limiting substrate eventually plateaued, suggesting that as with the minimalistic model, the mechanistic model can saturate.

In 1-D minimalistic models, saturation leads to a flattening of the concentration profile of activator in the peak, so that the maximum concentration stops increasing as the total protein amount rises (Fig. 1D). In the 2-D mechanistic model, the maximum concentration of each activator behaved differently depending on which protein was added to the system (Fig. 1I,J). When cytoplasmic Bem1-GEF substrate was limiting, the maximum Bem1-GEF-Cdc42 concentration in the peak fell while the maximum GTP-Cdc42 concentration rose (Fig. 1I). Conversely, when cytoplasmic GDP-Cdc42 was limiting, the maximum GTP-Cdc42 concentration in the peak fell while the maximum Bem1-GEF-Cdc42 concentration rose (Fig. 1J). Note that the total amount of both GTP-Cdc42 and Bem1-GEF in the peak always increased with increasing protein content, but the activator corresponding to the limiting substrate spread out more (the profiles shown are sections through a disc-shaped peak, so an apparently small increase in the activator concentration at the periphery of the peak actually represents a large increase in the total content of activator, offsetting the decreased concentration in the middle of the peak). Thus, concentration profiles for limiting species show saturation but those for non-limiting species do not.

We next asked whether saturation leads to co-existence in the mechanistic model. When simulations were initiated with two unequal peaks, competition occurred rapidly with low protein amounts, but slowed in a non-linear manner as the amount of protein increased (Fig. 1K), generating competition times that would be long compared to the yeast bud emergence timescale. In summary, saturation of polarity peaks is also evident in a mechanistic model with multiple species, and as in minimalistic models, the approach to saturation slows competition, driving the system towards coexistence.

### Equalization behavior in more complex models

In addition to positive feedback, the budding yeast Cdc42 system displays negative feedback (Howell et al., 2012; Kuo et al., 2014; Okada et al., 2013). Simulations of a mechanistic model with negative feedback modeled via inhibition of a Bem1-GEF-mediated positive feedback pathway were the first to show an unexpected novel behavior, in which starting unequal clusters neither competed nor coexisted, but instead equalized (Howell et al., 2012). An intuitive explanation for such equalization is that larger peaks are penalized by generating more negative feedback, allowing smaller peaks to compete successfully (Jacobs et al., 2019). However, equalization behavior does not necessarily follow from the presence of negative feedback (Chiou et al., 2018; Jacobs et al., 2019), suggesting that this intuition is insufficient to account for equalization. Here we seek to understand the features of polarity models that enable equalization.

Addition of a negative feedback loop to a minimalistic model (Fig. 2Ai) did not enable equalization (Chiou et al., 2018). However, in addition to the original report (Howell et al., 2012), addition of a negative feedback via activation of a Cdc42 GAP (Fig. 2Aii) (Jacobs et al., 2019) did enable equalization. In yeast cells, a major negative feedback pathway involves multi-site inhibitory phosphorylation of the GEF by Cdc42 effector PAKs (Kuo et al., 2014). We found that a mechanistic model incorporating this feedback loop (see methods: Fig. 2Aiii) also had the capacity to exhibit equalization (Fig. 2B). This suggested that some feature(s) absent from the first model (Fig. 2Ai) but shared by the others (Fig. 2Aii-iii) might explain equalization. One such feature is the addition of a new species (the GAP in model ii and the phosphorylated Bem1-GEF in model iii).

**Figure 2.**
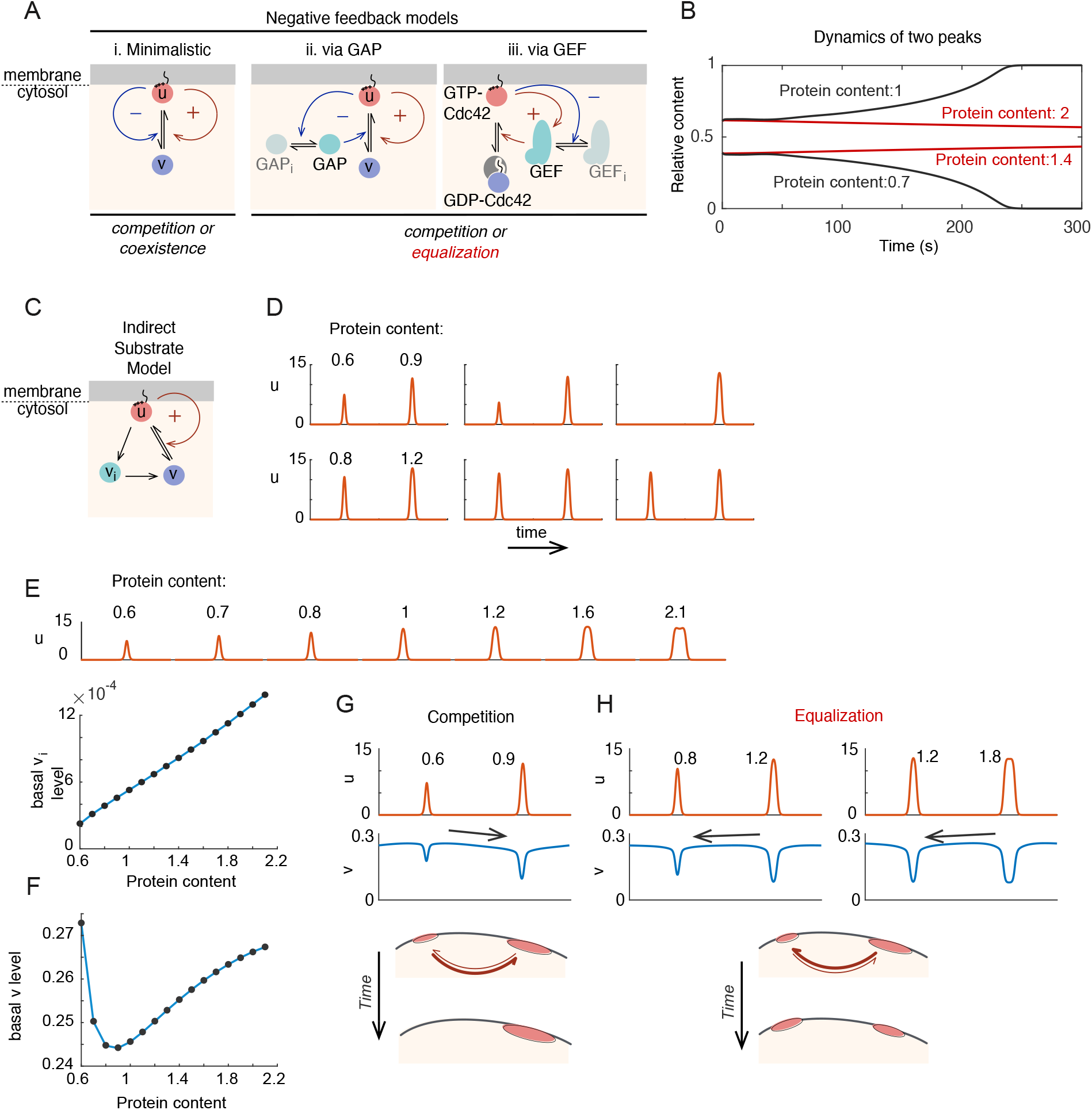
Basis for Equalization in more complex models. A) Schematic of models incorporating negative feedback. (i) In the minimalistic model, negative feedback adds a term in which *u* promotes conversion of *u* to *v* in a non-linear manner. (ii) This model incorporates conversion of an inactive GAP (GAPi) to an active GAP in a manner stimulated by *u*. The GAP promotes conversion of *u* to *v*, providing negative feedback. (iii) Simplified scheme of a mechanistic model where positive feedback occurs via recruitment of a GEF (Bem1-GEF in Fig. 1F) to sites containing GTP-Cdc42. Negative feedback occurs because GTP-Cdc42 promotes phosphorylation of the GEF, generating inactive GEFi. For details of the full model see Fig. 7F and Methods. B) With the GEF negative feedback model, two peaks can result in competition (protein content 0.7x, 1x) or equalization (protein content 1.4x, 2x) with varying Cdc42 content. C) Schematic of the indirect substrate model. In addition to the reactions from minimalistic MCAS model in Figure 1C, *u* can be converted into the indirect cytoplasmic substrate *v_i_*, which itself can convert to *v*. D) Unequal peaks in the indirect substrate model can yield competition or equalization. E) The basal level of indirect substrate increases with increasing protein content. Top row shows peak profiles at steady state for different starting protein content; graph shows corresponding steady state basal level of vi in the cytoplasm. F) The basal substrate level (*v*) first decreases but then increases with increased protein content. G) Competition occurs when the larger peak depletes total substrate more than the smaller. Protein content 0.6x, 0.9x. H) Equalization occurs when the smaller peak depletes total substrate more than the larger. Protein content 0.8x, 1.2x and 1.2x, 1.8x.

The GAP and the phosphorylated Bem1-GEF are neither substrates nor activators, and appear to play different roles in the polarity circuit. However, we noticed that they both provide a source of substrate: the GAP converts local GTP-Cdc42 into the substrate GDP-Cdc42, while the phosphorylated Bem1-GEF turns into the substrate Bem1-GEF upon dephosphorylation. Thus, in both cases a new species produced by the activator is highly mobile and generates a substrate in the cytoplasm. We reasoned that a larger peak of activator would generate more of this new species (GAP or phosphorylated Bem1-GEF) in its vicinity, and by generating more substrate this new species might reverse the concentration gradient of substrate in the cytoplasm, driving a flux of substrate towards the smaller peak to yield equalization.

Interestingly, the key to equalization in the hypothesis proposed above is not negative feedback per se, but rather the existence of a new high-mobility species created by an activator that can generate a substrate. To test our hypothesis, we modified the minimalistic model to include an “indirect substrate” species (Fig. 2C): In addition to direct conversion of the activator *u* to the substrate *v*, *u* can also convert to the indirect substrate *v_i_* in the cytoplasm, which then converts to *v*. We made the *u⟶ vi* reaction linear with *u*, such that this model lacks the non-linear negative feedback present in the mechanistic models discussed above, allowing us to probe whether equalization arises due to the presence of indirect substrate even without such negative feedback. This new model recapitulated the switch from competition to equalization behavior of the mechanistic models as the total protein amount in the system was increased (Fig. 2D). Previously, in the minimalistic model (Fig. 1C,D), basal levels of the cytoplasmic substrate *v* decreased until they reached a limit as peaks became larger. However, in this new model, basal levels of the indirect substrate *v_i_* rose steadily with peak size (Fig. 2E). The level of substrate *v* initially decreased but then rose as protein content was increased (Fig. 2F), presumably due to flux from *v_i_*. When two unequal peaks of activator were introduced, we found that whether the system showed competition or equalization was correlated with the relative basal levels of cytoplasmic substrate associated with each peak (Fig. 2G,H). When the larger peak was associated with lower substrate levels, the system displayed competition; when the smaller peak was associated with lower substrate levels, the system displayed equalization. Thus, a sufficient local production of indirect substrate by the larger peak can drive a flux of substrate towards the smaller peak, yielding equalization.

### Testing model predictions: multi-polar growth in large yeast cells

A simple prediction emerging from the polarity models discussed above is that cells should switch from competition to either coexistence or equalization as the amount of polarity “activators” in the clusters are increased. However, raising protein concentrations further in MCAS models generally drives them into a regime where Cdc42 activation would occur over the entire surface, yielding a uniform (depolarized) steady state. Thus, overexpression might lead to either multi-polarity or depolarization. Previous work indicated that overexpression of Cdc42 did not affect unipolar outcomes (Freisinger et al., 2013; Howell et al., 2012), while overexpression of Bem1 led to a small fraction of bipolar outcomes (Howell et al., 2009). These findings may suggest that some other component (e.g. the GEF) is limiting. Overexpression of the GEF, or a fusion protein between a PAK and the GEF, led to bipolar outcomes as well as depolarized cells, with depolarization becoming dominant upon co-overexpression of Cdc42 (Howell et al., 2012; Ziman and Johnson, 1994). The failure to observe frequent multi-budded cells upon overexpression may be due to excess protein concentrations that drive the system into the uniformly-high GTP-Cdc42 regime.

A different way to generate polarity domains with higher protein content in the models would be to increase the size of the modeled domain while keeping overall protein concentrations constant (Chiou et al., 2018): this should lead to multi-budded cells while avoiding depolarized outcomes. One way to increase cell size is to arrest the cell cycle, but in yeast this approach leads to cytoplasm dilution as biosynthesis fails to keep pace with volume growth (Neurohr et al., 2019). Instead, we utilized cytokinesis-defective yeast mutants to obtain large connected cells that continue cycling and presumably retain a normal overall protein composition.

The temperature-sensitive septin mutant *cdc12-6* is defective in cytokinesis at 37°C, generating chains of elongated, connected cells (Fig. 3A)(Hartwell, 1971). In the first cell cycle after switching to 37°C, all cells generated a single bud, which remained connected to the mother. In the second cell cycle, most cells formed a single bud despite having two cell bodies and two nuclei. This is consistent with the idea that these larger cells retained an effective competition mechanism that yields only a single winning polarity site to become the bud. However, some cells generated two buds simultaneously. The fraction of cells that are multi-polar increased further in the third and the fourth cell cycles (Fig. 3B). Similar multi-budded outcomes were observed in a conditional cytokinesis-defective *iqg1* strain (Fig. 3-fig. supp. 1a)(Shannon and Li, 1999), indicating that the phenotype is not specific to septin mutants. These results are consistent with the hypothesis that cells with higher total protein content (due to having a larger volume) can trigger a transition from competition to coexistence or equalization.

**Figure 3.**
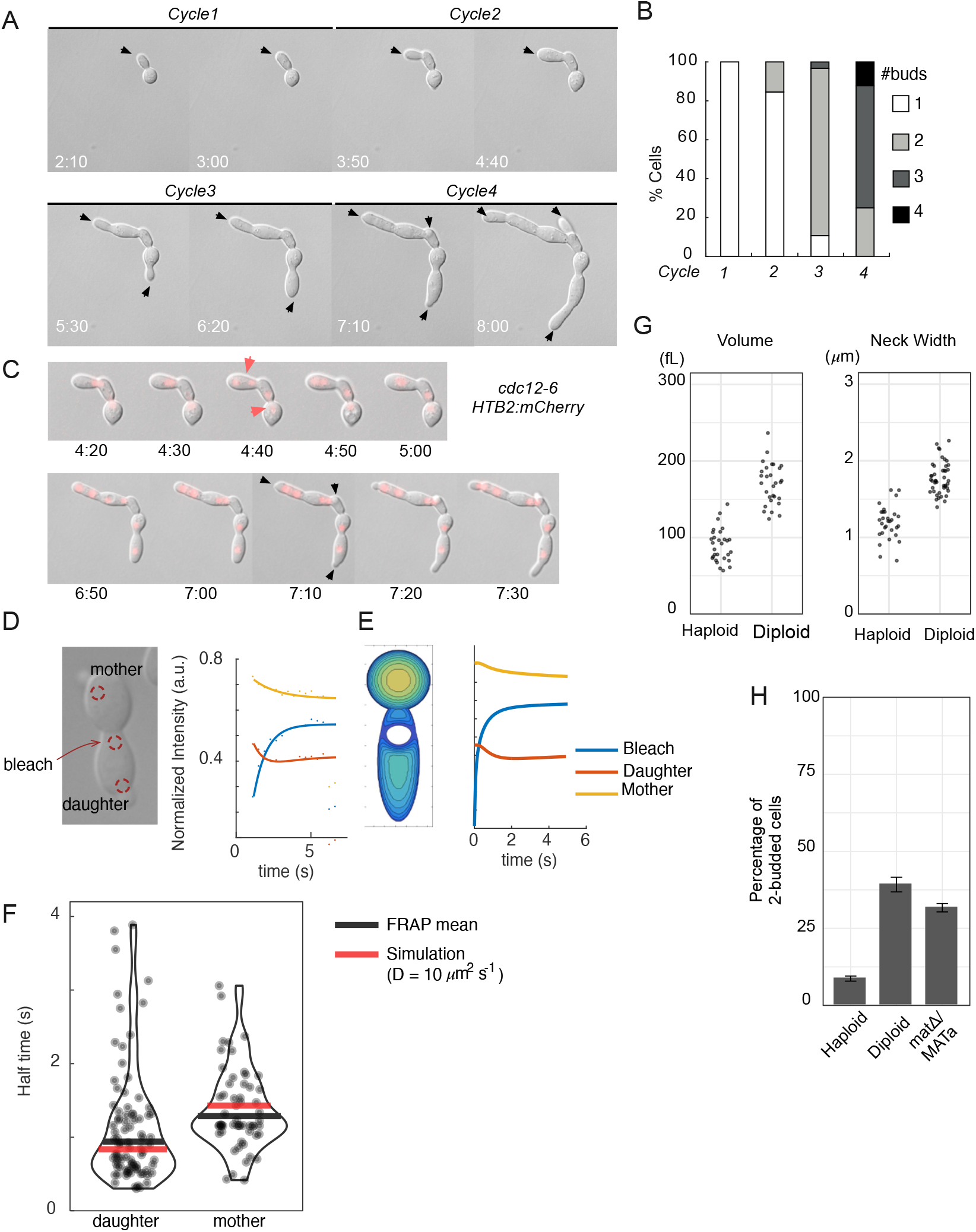
Large yeast cells can generate multiple buds. A) DIC time lapse movie of a cytokinesis-defective *cdc12-6* mutant (DLY20240) over 4 budding cycles at restrictive temperature (37°C). Black arrows indicate growing buds. Time in h:min. B) The number of buds generated in each cell cycle was scored for N = 35 cells. C) The nuclear probe Htb2-mcherry exhibits roughly synchronous nuclear divisions (top, red arrows) and bud emergence (bottom, black arrows) indicating that the entire cell remains connected. Time in h:min. D) Effect of neck on cytoplasmic diffusion. Cells expressing cytoplasmic GFP (DLY22957) were bleached at a spot in the daughter compartment immediately adjacent to the neck. Dynamics of fluorescence intensity in each cell were measured in the mother and daughter compartments equidistal to the bleach site (red circles), and fitted to exponential curves. E) Simulated FRAP setup (left) for the same experiment yielded similar recovery curves (right). F) Half-times extracted from exponential decay curves from FRAP experiments (each dot is one FRAP experiment, black line: median) and simulation data (red line). G) Volume and neck width of diploid (DLY20569) and haploid (DLY9455) *cdc12-6* cells in the second cell cycle at restrictive temperature. H) Percentage of 2-budded cells observed in the second cell cycle at restrictive temperature for diploid (DLY20569) and haploid (DLY9455) *cdc12-6* cells. *mat*Δ/MATa cells (DLY22887) are diploid but have a haploid mating type.

**Figure 3-supporting Figure 1.**
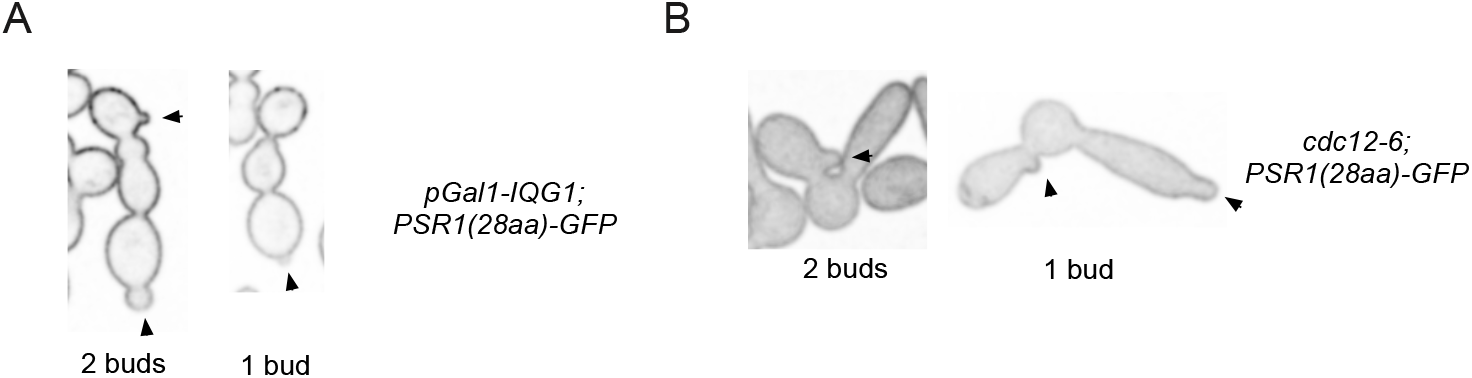
A) Medial-plane images of 1- and 2-budded cells after *IQG1* shut-off for X h (DLY22875). B) Medial-plane images of 1- and 2-budded cells after shift of *cdc12-6* cells (DLY22915) to restrictive temperature for X h. Membrane probe Psr1-GFP. Arrowheads indicate buds.

A potential caveat to our conclusion is that in later cell cycles, the mutant cells might complete some form of cell division, so that the cytoplasm was not truly connected in the apparent multi-budded cells. Arguing against this possibility, budding and nuclear division were roughly synchronous within a chain of cell compartments (Fig. 3C), suggesting cytoplasmic communication between compartments. More definitively, the continuity of the cytoplasm was confirmed by using the plasma membrane probe Psr1^1-28^-mCherry (Fig. 3-fig. supp. 1b)(Siniossoglou et al., 2000).

Even with a continuous cytoplasm, it was possible that the narrow necks between compartments could impede cytosolic communication (diffusion), preventing competition across compartments. To test this possibility, we conducted photo-bleaching experiments on cycle 2 *cdc12-6* cells expressing free cytosolic GFP. We bleached a spot in the daughter compartment and monitored the effect on fluorescence in the same or the neighboring mother compartment (Fig. 3D, F). Fluorescence intensity time-courses were fit to exponential curves, allowing us to extract characteristic diffusion times across the same distance with or without a neck in between. We found that communication by diffusion was very rapid, and the presence of the neck slowed communication only very mildly (Fig. 3D). This small effect can be explained simply by the neck geometry, as *in silico* simulated photo-bleaching of a system with similar geometry produced a comparable effect (Fig. 3E,F). Thus, the neck does not appear to impose a diffusion barrier beyond that expected for the narrowing of the isthmus.

A remaining caveat is that the small effect on cytoplasmic diffusion imposed by the neck geometry might suffice to yield multi-polarity. To distinguish whether increased volume or neck geometry is the dominant contributing factor for multipolar outcomes, we compared haploid and diploid mutant cells. Diploid cells have larger volume (which correlates with total protein content) but also wider necks (which would provide a smaller impediment to diffusion) compared to haploids (Fig. 3G). Despite having wider necks, the larger diploids generated more 2-budded cells than did the haploids (Fig. 3H). This was due to cell size and not mating type, as MATΔ/MAT**a** cells (cells that have the large size of a diploid but the mating type of a haploid) behaved similarly to normal diploids (Fig. 3H). Taken together, our findings indicate that the large cells generated following failure of cytokinesis can yield multi-budded outcomes, and that such outcomes can be predominantly attributed to the larger cell size.

### Polarity dynamics in cytokinesis-defective cells

Larger cell size could lead to a multi-polar outcomes by affecting two aspects of polarity dynamics: Having more polarity patches that initially form during polarity establishment, or changing the subsequent behavior of the polarity system (competition, co-existence, or equalization). To evaluate these features, we introduced polarity probes into *cdc12-6* cells. We used Bem1-tdTomato as a polarity probe (Howell et al., 2012), and employed a *cdc12-6 rsr1*Δ genetic background, to avoid complexities associated with Rsr1-mediated bud-site-selection, which biases the location of polarity patches and slows competition by unknown mechanisms (Bi and Park, 2012; Wu et al., 2013).

In wild-type (not cytokinesis-defective) cells, Bem1 and Cdc42 localize to the mother-bud neck during cytokinesis and polarize at presumptive bud sites during late G1. The neck-localized Cdc42 is thought to be predominantly inactive during cytokinesis (Atkins et al., 2013), and several effectors of Cdc42 (including the PAK Cla4) do not co-localize with Cdc42 and Bem1 at the neck (Holly and Blumer, 1999). In *cdc12-6* cells, we expected that we would not see the neck localization (as these mutants do not undergo cytokinesis), and only detect polarization in late G1. However, surprisingly Bem1 manifested two rounds of localization. To understand the timing of these events, we used Whi5-GFP as an indicator of cell cycle stage. Whi5 localizes to the nucleus from late mitosis until early G1, but is cytoplasmic from late G1 through metaphase (Costanzo et al., 2004; de Bruin et al., 2004; Skotheim et al., 2008). The first round of Bem1 localization in *cdc12-6* cells (Fig. 4A, blue arrow) occurred after Whi5 entered the nuclei (Fig. 4A, green arrow; i.e. at the normal time of cytokinesis), and the second round occurred after Whi5 exited the nuclei (Fig. 4A, green arrow; i.e. at the normal time of polarization)(Fig. 4A, red arrows). Given its timing, the first round of localization likely represents a failed attempt at cytokinesis. Consistent with that view, in rare cases where cytokinesis did succeed, the cytokinesis sites coincided with sites of first-round Bem1 localization (Fig. 4-fig. supp. 1a). Moreover, the effector Cla4 did not co-localize with Bem1 at the first-round locations, and only co-localized with Bem1 in the second round of polarization (Fig. 4-fig. supp. 1b). These findings indicate that *cdc12-6* cells make an unexpected attempt at cytokinesis, generally at sites away from the neck. Nevertheless, we can monitor polarization dynamics by focusing on the second round of Bem1 polarization.

**Figure 4.**
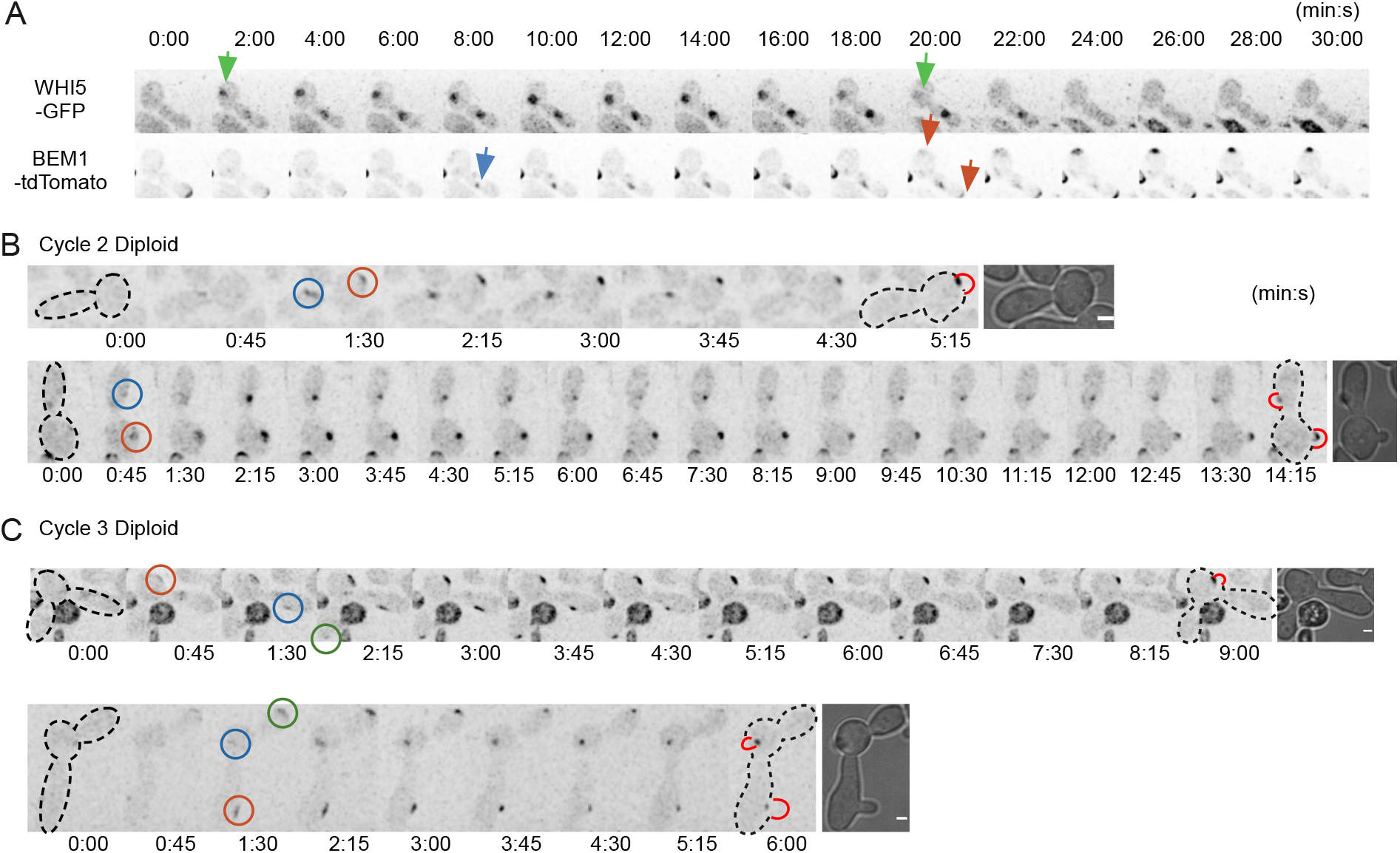
Polarity patch behaviors in large *cdc12-6* cells. A) Bem1 clusters twice, during aborted cytokinesis and polarization. Time-lapse imaging of cell cycle (Whi5-GFP) and polarity (Bem1-tdTomato) probes in *cdc12-6* cells at restrictive temperature (DLY22920). Green arrows indicate Whi5 nuclear entry (marking mitotic exit) and Whi5 nuclear exit (marking start of the next cell cycle). Blue arrow indicates the clustering of Bem1-tdTomato before start, indicative of attempted cytokinesis. Red arrows indicate polarization of Bem1-tdTomato after start. Time in min:s. B) Cells that start with 2 polarity patches can show competition or coexistence. Example cells demonstrating typical Bem1 behaviors in *cdc12-6* diploids (DLY15376) in the second G1 phase after switching to restrictive temperature. Circles indicate initial polarity sites. Red bulges indicate buds. Top: competition yields one bud. Bottom: coexistence yields two buds. C) Example cell from the third G1 phase after switching to restrictive temperature. This cell initially shows 3 polarity patches but makes 2 buds. Scale bar = 2 μm.

**Figure 4-supporting Figure 1.**
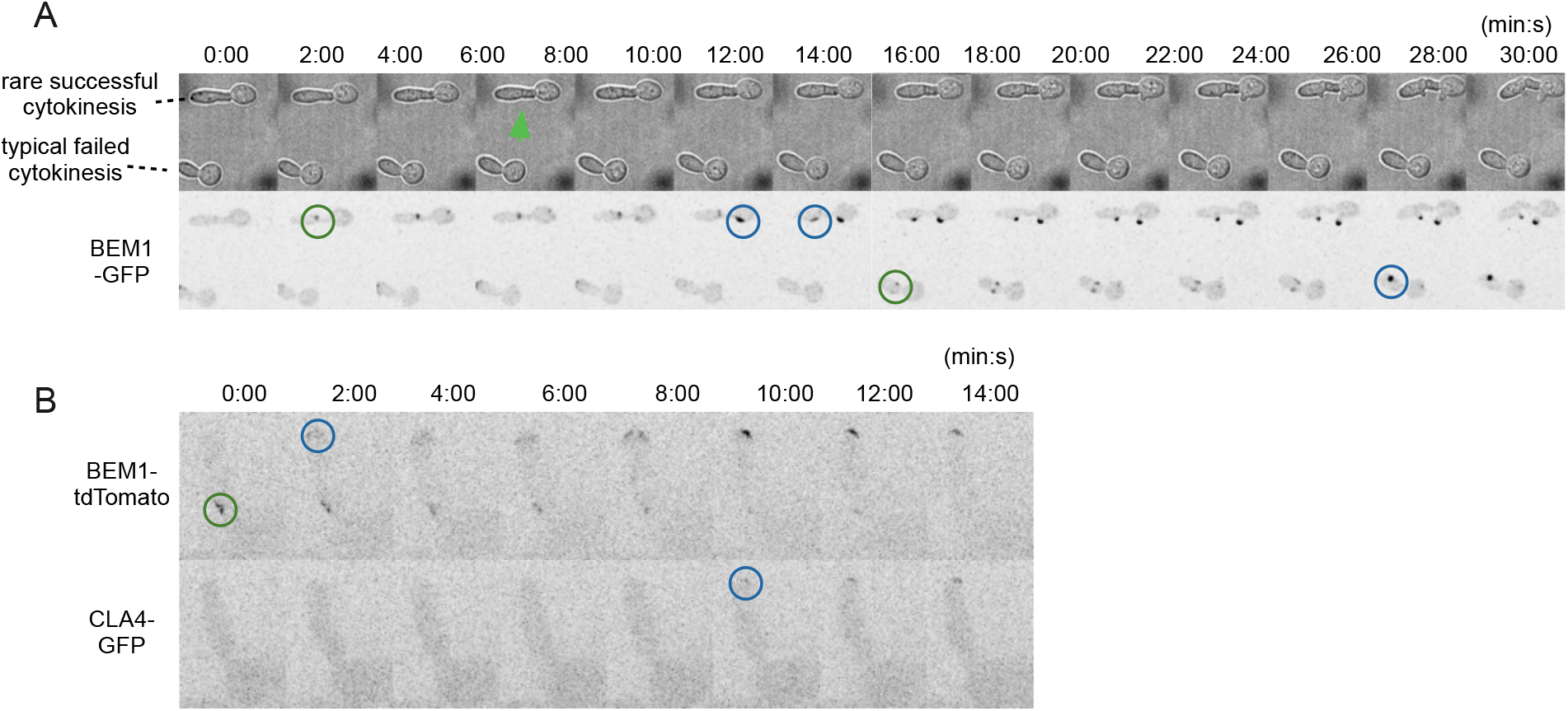
A) A rare successful cytokinesis in *cdc12-6* cells (DLY16767) at restrictive temperature. Bem1-GFP localized to cytokinesis site (green circle) and then polarity site (blue circle). B) Cla4 does not localize to cytokinesis site. Bem1-tdTomato localized to cytokinesis site (green circle) and then polarity site, while Cla4-GFP only localized to the polarity site (DLY23359).

As cells entered the second cell cycle after switching to 37°C, Bem1 became concentrated at one or more initial polarity patches (Fig. 4B). Some of these initial patches then disappeared, presumably as a result of competition. Such competition could be seen within individual cell compartments or between cell compartments connected by necks. However, in some cases the initial patches persisted and gave rise to two buds. As cells entered the third cell cycle, Bem1 sometimes localized to 3 patches. The outcome in these cases was variable, with competition leaving one or two patches that gave rise to buds (Fig. 4C). Thus, unipolar versus multipolar outcomes appear to depend both on the number of initial patches that form and on whether they are subsequently eliminated.

To assess how cell size might affect initial patch numbers, we compared *cdc12-6 rsr1*Δ cells in the second cell cycle versus the third cell cycle after switching to 37°C, as well as haploid and diploid mutant cells. As expected, third-cycle cells were larger than second-cycle cells, and diploids were larger than haploids (Fig. 5A). The number of initial patches formed by each cell increased in a manner correlated with cell length (Fig. 5B), suggesting that cell length can influence initial polarization. However, the locations at which patches formed were non-random, with a preference for bud tips and mother cell locations (Fig. 5C), suggesting that this may not be a simple case of symmetry breaking. When cells formed two initial patches, the distance between the patches was highly variable (Fig. 5D). This is contrary to the expectation for classical Turing-type models, which tend to form patches separated by a characteristic length scale (Meinhardt, 2008). However, it is consistent with predictions from MCAS models (Goryachev and Pokhilko, 2008). Of the cells that formed 2 initial patches, the fraction that yielded 2-budded cells also increased with cell size (Fig. 5E). Thus, the probability of successful competition between patches decreases as cells grow larger, consistent with a transition from competition to co-existence or equalization regimes.

**Figure 5.**
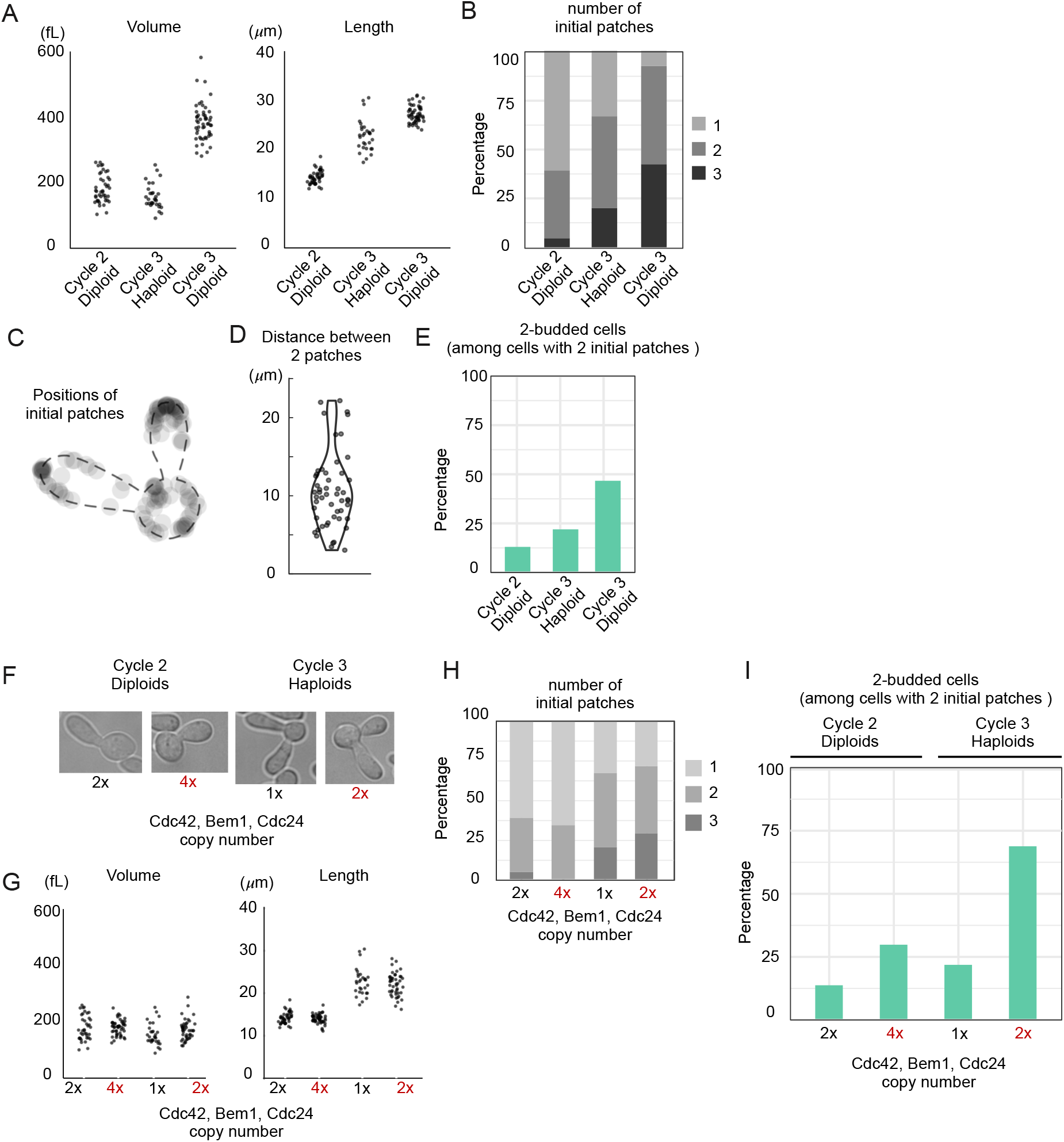
Increasing protein content promotes switch from unipolar to multipolar outcomes. A) Cell volume and length in three populations of *rsr1Δ cdc12-6* cells at restrictive temperature: diploids (DLY15376) in the second or third cell cycle and haploids (DLY9453) in the third cell cycle. B) The number of initial polarity patches in each population. C) The distance between 2 patches in diploids and haploid cells exhibits a broad distribution. D) Initial clusters form at non-random positions. E) Larger cells show more frequent multipolar outcomes. Percentage of of 2-budded cells (bipolar outcomes) among those that established 2 initial patches. F) Examples of diploid (DLY23308) and haploid (DLY23302) *rsr1Δ cdc12-6* cells with double the gene dosage of *CDC42*, *BEM1* and *CDC24* were compared to cells with a normal dose (DLY15376, DLY9453). G) Cell size is unaffected by polarity gene dosage (strains as in F). H) The number of initial patches does not vary systematically as a function of polarity gene dosage (strains as in F). I) Similar sized cells with more polarity proteins show more frequent multipolar outcomes. The percentage of 2-budded cells within the subpopulation that established 2 initial patches (strains as in F).

In MCAS models, the main effect of larger size on system behavior is due to the increased total abundance of polarity proteins in the system, rather than the increased length or volume per se (Chiou et al., 2018). To ask whether this is also the case in our mutants, we integrated an additional copy of the *CDC42*, *CDC24* (GEF), and *BEM1* genes into our mutant cells (Fig. 5F). This did not affect the volume or length of the cells (Fig. 5G), and there was no systematic effect on the number of initial patches formed (Fig. 5H). However, among the cells that established 2 initial patches, the frequency of multi-polar outcomes increased (Fig. 5I). We conclude that increasing polarity protein content is sufficient to decrease the effectiveness of competition, yielding multi-polar outcomes.

### Are multi-polar outcomes a result of equalization or co-existence?

In MCAS models, multipolar outcomes arose in two ways: through equalization (where a smaller cluster grows faster than a larger one) or saturation (where competition between clusters slows dramatically). As discussed above, equalization behavior can arise from the presence of an indirect substrate species, a realization motivated by the observation that a phosphorylated and inhibited form of the GEF Cdc24 (which behaves as an indirect substrate) accumulates during polarization in yeast cells. To ask whether multipolar outcomes in our cytokinesis-defective yeast cells were due to this hypothesized indirect substrate, we constructed a mutant strain in which the wild-type Cdc24 was replaced with a non-phosphorylatable version, Cdc24^38A^. We found that this did not diminish multipolar outcomes. Indeed, this strain made two-budded cells at a higher frequency than size-matched Cdc24-wild-type controls (65% vs 12% multipolar for Cdc24^38A^ vs Cdc24 second-cycle cells, respectively). This mutant is complicated to interpret because in addition to eliminating the “indirect substrate” phosphorylated Cdc24 species, it increases the amount of active Cdc42 in the polarity patches (Kuo et al., 2014). Increased active Cdc42 would be predicted to drive the system towards saturation, potentially explaining the increase in multipolar outcomes.

Although phosphorylated Cdc24 is not required for multipolar outcomes, it remained possible that some other species (e.g. a mobile GAP or another indirect substrate) might be causing equalization. Equalization and co-existence behaviors make different predictions about how a hypothetical starting cell with two unequal polarity patches would evolve. In an equalization scenario, the initially unequal patches should evolve towards two equal patches. In a saturation scenario, on the other hand, the unequal patches could co-exist while remaining unequal. And if the cell is approaching but has not reached saturation, then the unequal patches would slowly diverge in protein content as competition proceeds. We therefore asked which of these scenarios best accounts for the behavior of cytokinesis-defective yeast cells.

A difficulty in distinguishing equalization and competition behaviors in cells with wild-type Cdc24 is that negative feedback through Cdc24 phosphorylation can cause oscillation in the protein content of the polarity patches, obscuring the underlying processes (Howell et al., 2012; Kuo et al., 2014). The *CDC24*^38A^ strain short-circuits this negative feedback and does not show oscillations, allowing us to circumvent this complexity. Focusing on *CDC24*^38A^ cells that had two starting patches in the second cycle at 37°C, we tracked the total Bem1 fluorescence in each patch over time until around the time of bud emergence. We quantified the fluorescence in each patch as a fraction of the sum total in both patches, for each timepoint. Time-courses for 20 cells are shown in Fig. 6. These cells exhibit a continuum of behaviors that can be classified as competition (10 cells: the larger patch grows and the smaller one shrinks), coexistence (9 cells: the relative Bem1 content in the patches stays approximately constant), or equalization (1 cell: the larger patch shrinks and the smaller one grows). Thus, most cells demonstrate either competition (e.g. Fig. 6A, cell iv) or coexistence (e.g. Fig. 6A, cell xiii). Interestingly, 3 of the cells with the slowest competition (Fig. 6A, cells viii-x) had not completed competition by the time of bud emergence, and went on to grow two buds (e.g. Fig. 6A, cell viii). We call this phenotype “aborted competition”.

**Figure 6.**
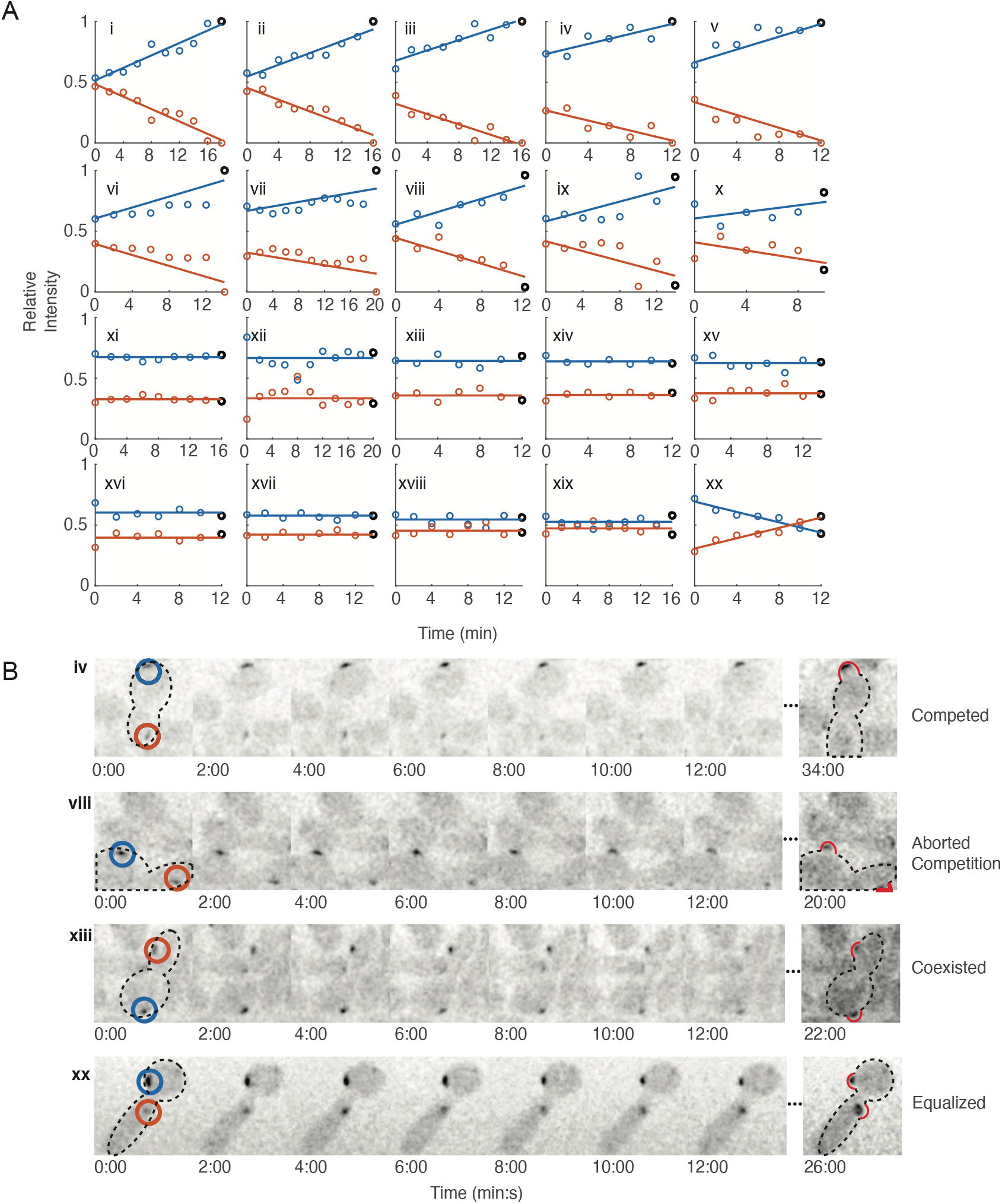
Competition, Coexistence, and Equalization in yeast. A) Relative amounts of Bem1 over time were quantified from Bem1-GEF sum intensity traces from z-stacks of 20 *rsr1Δ CDC24^38A^* cells (DLY21100) that had 2 initial polarity patches. Patches that eventually led to bud-emergence are indicated by black circles at the last time point. Cells i-vii: competition (1 bud). Cells viii-x: aborted competition (2 buds). Cells xi-xix: coexistence (2 buds). Cell xx: equalization (2 buds). B) Example traces that fall into each category: Roman numerals refer to panels in A). Circles highlight initial patches. Red bulges in the last panels indicate buds.

If competition between two polarity patches occurs up until bud emergence, why would competition then stop and allow both buds to grow continuously? This behavior suggests that after bud emergence the two buds become somehow insulated from each other in terms of polarity behavior, even though FRAP experiments confirm that their cytoplasms remain connected (Fig. 3D-F). The switch from competition to coexistence/equalization may indicate that some aspect of the polarity circuit changes at the time of bud emergence.

## Discussion

### Increasing cell size promotes a switch from uni-to multi-polarity

Using conditional cytokinesis-defective yeast mutants, we found that larger cells produced progressively higher numbers of buds. Control experiments indicated that the undivided cell shared a single cytoplasm with negligible diffusion barriers. Increasing numbers of polarity clusters with cell size is predicted by three different classes of simple mathematical pattern-formation models relevant to polarity establishment. Each class of models provides a different explanation for the switch from uni-polar to multi-polar outcomes.

Classical (not mass-conserved) Turing models exhibit a characteristic length scale, such that clusters of an activator form spontaneously at spatial intervals corresponding to this length scale. Cells develop one or more clusters depending on how large the cell is compared to the characteristic length scale (Meinhardt, 2008).

Mass-conserved Activator-Substrate (MCAS) models can develop variable numbers of initial activator clusters, but as substrate is depleted from the cytoplasm the clusters compete with each other. Eventually, competition leads to a single-cluster steady state, but the timescale of competition slows dramatically as the protein content in the system is increased (Chiou et al., 2018; Ishihara et al., 2007). Cells with a small activator/substrate content exhibit rapid competition and evolve to have a single cluster, whereas those with larger activator/substrate content maintain two (or more) clusters on biologically relevant timescales.

More complex mass-conserved models incorporating more polarity regulators can switch from a regime exhibiting competition (where a larger cluster grows faster than a smaller cluster) to a regime exhibiting equalization (where a smaller cluster grows faster than a larger cluster) as parameters change (Howell et al., 2012). The basis for this behavior is discussed in more detail below. For now, we note that a switch from competition to equalization as cells become larger could also explain the observed switch from uni-polar to multi-polar outcomes.

The number of initial polarity clusters in our cells increased with increasing cell length, consistent with Turing models in general. However, there was no obvious preferred length scale for the distance between clusters, as would be expected from classical Turing systems. A potential explanation for the increase in initial clusters stems from the observation that cluster locations were non-random, with a preference for cell tips. For the geometry of our cytokinesis-defective cells, the number of tips correlates with cell length, because longer cells arise from the formation of additional buds. Thus, the number of initial polarity sites may reflect the specifics of our experimental system rather than a general feature of the polarity circuit.

Those cells that did form more than one initial polarity cluster often exhibited competition between clusters. Competition emerges as a consequence of mass conservation, and suggests that MCAS models provide the explanation that best fits the behavior of the yeast system. Also consistent with MCAS models, analyses of cells with different ploidy suggested that a switch from competition to coexistence scaled with cell volume (and hence protein content) rather than cell length. Moreover, additional copies of genes encoding polarity proteins led to a marked increase in the frequency of multi-polar outcomes with no change in cell size. These findings strongly support the idea that multi-polar outcomes arise due to a change in the timescale of competition (or equalization: see below).

### Competition, saturation, and aborted competition

Analyses of minimalistic MCAS models demonstrated that competition between activator clusters was inevitable, and that the timescale of competition was determined by a single dominant factor, which we refer to as saturation (Chiou et al., 2018; Jacobs et al., 2019; Otsuji et al., 2010). When activator concentration in two or more clusters approaches a saturation point set by system parameters, competition slows dramatically, allowing co-existence of the clusters on biologically relevant timescales. We tested whether these findings from minimalistic one-activator, one-substrate models would hold in a more complex and realistic model of the yeast polarity circuit, with two activators, two substrates, and two other species. With some additional complexity discussed below, our findings support the idea that saturation can also account for the behavior of more complex models.

In minimalistic MCAS models, positive feedback ensures that addition of more substrate/activator to the system results in conversion of more substrate to activator. Due to positive feedback, the concentration of substrate is depleted below the level obtained with less substrate/activator in the system at steady state. In addition, the concentration profile of activator in the cluster changes in a characteristic way as the local activator concentration approaches the saturation point, flattening from a sharp peak to a mesa. In the multi-component model, both substrate depletion and the activator concentration profile are dependent on the specific species. Depending on the relative amounts of the different species, different substrates may become limiting. For the limiting species, addition of more protein will generally lead to depletion of the cytoplasmic substrate, similar to saturation in the minimalistic models. However, other (non-limiting) species can display increasing cytoplasmic substrate concentration as more protein is added. Moreover, addition of one species can lead to a switch in the identity of the limiting species.

As with minimalistic MCAS models, the timescale of competition in the mechanistic multi-component model slows as more protein is added to the system, in a manner consistent with saturation of the limiting species. In principle, this could explain why larger yeast cells can make more than one bud. Tracking a polarity marker in time-lapse imaging, we found two predominant behaviors among large yeast cells that generated two initial polarity clusters: one subset exhibited competition between clusters to yield a single bud, while the other exhibited apparent coexistence between clusters yielding two buds. These behaviors are consistent with a scenario in which competition occurs on timescales that vary in individual cells depending on the degree of saturation of the limiting species in that cell. As cells become larger, they are more likely to approach saturation, slowing competition to the point that two clusters are present at the time of bud emergence, yielding two buds.

A smaller group of cells exhibited competition that failed to go to completion, so that the uneven clusters both generated buds. This behavior, which we call “aborted competition”, is at odds with the predictions of MCAS models, in which competition accelerates as the content of activator in the peaks becomes more uneven and always goes to completion. This discrepancy may indicate that some aspect of the polarity circuit changes at around the time of bud emergence, reducing the efficacy of competition. Cell cycle control by the cyclin/CDK system provides one plausible candidate regulator that could prompt such a change in polarity circuit behavior (Knaus et al., 2007; Moran et al., 2019; Sopko et al., 2007; Witte et al., 2017).

### Equalization

Unlike minimalistic MCAS models, more complex models (inspired by experimental observations that the yeast polarity circuit also exhibits negative feedback) can yield equalization of polarity peaks in some parameter regimes (Howell et al., 2012; Jacobs et al., 2019). The behaviors of all of the models considered in this paper are summarized in Fig. 7. Local negative feedback provides an intuitive rationale for equalization: by penalizing a larger peak more than a smaller peak, a localized negative feedback loop could switch the competitive advantage towards the smaller peak. However, previous work showed that incorporating negative feedback into minimalistic 2-component MCAS models did not lead to a switch from competition to equalization (Chiou et al., 2018), suggesting that this rationale is incorrect, or at least incomplete. Consistent with that conclusion, a recent analysis of MCAS models with negative feedback via activation of a GAP found that in order to yield equalization, the GAP must be more mobile than the GTP-Cdc42 (Jacobs et al., 2019). As higher mobility of the GAP would delocalize the negative feedback from the larger peak towards the smaller peak, this observation cannot easily be accommodated in a framework where equalization arises from localized negative feedback. Here we propose a novel mechanism for equalization that does not require negative feedback, but can account for the behavior of the more complex models that incorporate negative feedback.

**Figure 7.**
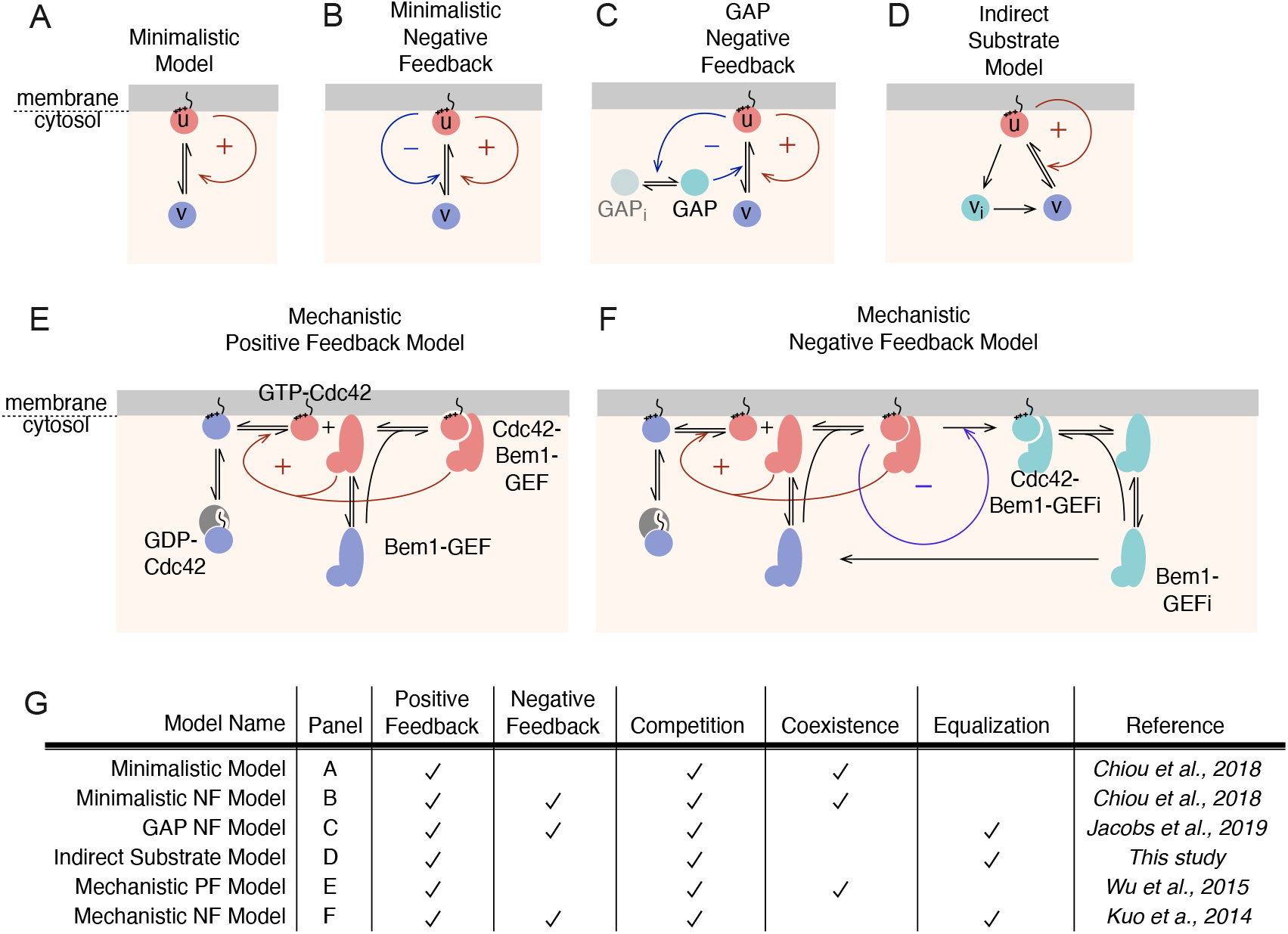
Summary of polarity model behaviors. A-D) Schematic of minimalistic MCAS models from Fig. 1 and Fig. 2. E,F) Schematic of mechanistic models without (E) or with (F) negative feedback (note that a simplified version of F was shown in Fig. 2Aiii). In addition to the reactions from the positive-feedback model in E, Bem1-GEF-Cdc42 complexes contain a kinase that phosphorylates the GEF in other Bem1-GEF-Cdc42 complexes (F, blue arrow). Phosphorylated species (indigo) lack GEF activity, but can still undergo reversible binding to Cdc42-GTP and exchange between membrane and cytoplasm. Dephosphorylation of inactive Bem1-GEF is assumed to occur only in the cytoplasm. G) Summary of model features and behaviors. Panel refers to schematics above. References describe first report of the models.

Just as competition between peaks involves a net flux of cytoplasmic substrate from the smaller peak to the larger peak, equalization must involve a net flux of cytoplasmic substrate from the larger peak to the smaller peak. As illustrated in Fig. 2H, the substrate concentration gradient that drives the net flux occurs between the outskirts of each peak, not directly below the peak where there is a local dip in substrate concentration. Thus, a key to equalization is the generation of more substrate at the outskirts of the larger peak than at the outskirts of the smaller peak. This rationale explains why GAPs that remain localized are ineffective at driving equalization: the substrate produced by the local GAP may help to fill the dip in cytoplasmic substrate directly beneath the peak but leaves the concentration at the outskirts unchanged. Endowing the GAP with a higher mobility allows it to generate new substrate more broadly, affecting the substrate concentration on the outskirts of each peak and reversing the substrate concentration gradient in the cytoplasm to promote equalization.

Our analyses show that equalization can also be observed in simple models that lack negative feedback (Fig. 7). The key to that behavior was the existence of an “indirect substrate” species: a cytoplasmic (high-mobility) species that cannot be directly be converted into an activator. The existence of such a species was inspired by the observation that in yeast, negative feedback occurs via phosphorylation of the Cdc42-directed GEF, yielding a cytoplasmic GEF incapable of activating Cdc42 until it had been dephosphorylated. Because the indirect substrate is produced by the activator and released into the cytoplasm, a larger activator peak releases more indirect substrate. As the indirect substrate (phosphorylated GEF) is converted to substrate (dephosphorylated GEF), it drives a flux of substrate from the larger to the smaller peak, feeding growth of the smaller peak to drive equalization. Even a system with very transient (rapidly dephosphorylated) indirect substrate could trigger a switch from competition to equalization. Notably, the existence of an indirect substrate species yields equalization even in the absence of negative feedback. As with GAP-containing models, the critical factor is that there be an activator-dependent mechanism for a peak to generate substrate over a broader area than that of the peak itself, so that substrate levels rise on the outskirts of the peak, reversing the substrate concentration gradient.

We note that what appears to be equalization can occur even without an indirect substrate in models that do not assume mass conservation (Jacobs et al., 2019). However, the basis for such equalization is mechanistically different. In mass-conserved models with an indirect substrate, equalization reflects a flux of cytoplasmic substrate from the larger peak to the peak. But in models that allow synthesis/degradation of polarity factors, each peak’s activator concentration profile reflects a local steady state, and the clusters are equal because they share the same synthesis and degradation parameters.

Does equalization account for the multi-polar outcomes we documented in large yeast cells? If phosphorylated GEF was important for such outcomes, then mutants with non-phosphorylatable GEF should exhibit a reduced frequency of multi-budded cells. Instead, they showed a higher frequency compared not normal-GEF cells. That does not eliminate the possibility that some other indirect substrate species or a high-mobility GAP may promote equalization. However, we only observed one instance in which an initially smaller peak appeared to grow at the expense of a larger peak, as predicted by equalization (Fig. 6). We tentatively conclude that for the cell populations we examined, multi-polar outcomes in the budding yeast are probably due to slow competition/coexistence rather than equalization.

### Implications for other systems

The *Saccharomyces* polarity circuit has presumably been evolutionarily selected to produce uni-polar outcomes, which are beneficial during budding and mating in this genus. However, this polarity circuit is highly conserved among ascomycetes that display other growth modes. *Schizosaccharomyces pombe* naturally switch from uni-polar to bi-polar growth during each cell cycle (Martin and Chang, 2005), *Ashbya gossypii* form branching hyphae that exhibit increasing number of polarity sites as each cell grows (Knechtle et al., 2003), and yeast cells of *Aureobasidium sp.* generate variable numbers of buds simultaneously (Mitchison-Field et al., 2019). This phenotypic diversity may be enabled by a polarity circuit that allows a switch between competition, coexistence, and equalization behaviors in response to appropriate tuning of parameter values. Similar principles may apply to other systems where activator species produce robustly tunable numbers of polarity sites.

## Methods

### Yeast strains

All yeast strains (Table S1) are in the YEF473 background (*his3-Δ200; leu2-Δ1; lys2-801amber; trp1-Δ63; ura3-52*)(Bi and Pringle, 1996). The *cdc12-6* mutation in the YEF473 background was a gift from John Pringle (Stanford University). The *rsr1* deletion (Schenkman et al., 2002), *GAL4BD-hER-VP16* construct (Takahashi and Pryciak, 2008), and *CDC24*^*38A*^ mutation (Kuo et al., 2014; Wai et al., 2009) were described previously, as was tagging at the endogenous loci for the fluorescent probes *BEM1-GFP* (Kozubowski et al., 2008), *BEM1-tdTomato* (Howell et al., 2012), *CDC3-mCherry* (Howell et al., 2009), *CLA4-GFP* (Wild et al., 2004), *HTB2-mCherry* and *WHI5-GFP* (Doncic et al., 2011). Standard yeast genetic crosses were used to generate all of the strains.

To generate a strain with regulatable expression of *IQG1*, the first 500 bp of the *IQG1* open reading frame were amplified by PCR and cloned downstream of the *GAL1* promoter in YIpG2 (Richardson et al., 1989) to generate DLB2126. Digestion at the unique *Nhe*I site targets integration of this construct at *IQG1*, making Iqg1 expression galactose-dependent and shut off on glucose media. To introduce a 3xHA epitope tag at the C-terminus of *CDC24*, we used a pFA6-series plasmid template and the PCR-based one-step replacement method (Longtine et al., 1998). To delete the MATα locus, we used a pFA6-series plasmid template and the PCR-based one-step replacement method (Longtine et al., 1998) to replace a part of the locus inactivating the divergent α1 and α2 genes while leaving the surrounding genes intact. In a haploid, this deletion converts an α mating type to an a mating type. In a diploid, this deletion converts the strain to a mating type.

To label the plasma membrane, we expressed a fusion between the N-terminal 28 residues of Psr1 and GFP. The Psr1 N-terminal fragment is myristoylated and doubly palmitoylated, targeting GFP to the plasma membrane (Siniossoglou et al., 2000). The construct was cloned between the *TEF1* promoter and *ADH1* terminator sequences in a pRS305 (Sikorski and Hieter, 1989) backbone, generating plasmid DLB4206. Digestion at the unique *Ppu*MI targets integration at the *LEU2* locus.

To express an extra copy of *CDC42*, the *CDC42* gene (open reading frame plus 500 bp upstream and 250 bp downstream) was cloned into the integrating plasmids pRS304 and pRS306 (Sikorski and Hieter, 1989), generating plasmids DLB3904 and DLB4115 respectively. Digestion of DLB3904 at the unique *Sty*I site was used to target integration of the *TRP1*-marked plasmid at *CDC42*, and digestion of DLB4115 at the unique *Stu*I site was used to target integration of the plasmid at *URA3*. To express an extra copy of *CDC24*, the *CDC24-3HA* gene (open reading frame plus upstream and downstream sequence) was cloned into the integrating plasmid pRS306 (Sikorski and Hieter, 1989), generating plasmid DLB4134. Digestion at the unique *Pst*I site was used to target integration of the plasmid at *URA3*. To express an extra copy of *BEM1-GFP*, the *BEM1-GFP* gene (open reading frame plus upstream and downstream sequence) was cloned into the integrating plasmid pRS304 (Sikorski and Hieter, 1989), generating plasmid DLB2997. Digestion at the unique *Bam*HI site was used to target integration of the plasmid at *BEM1*.

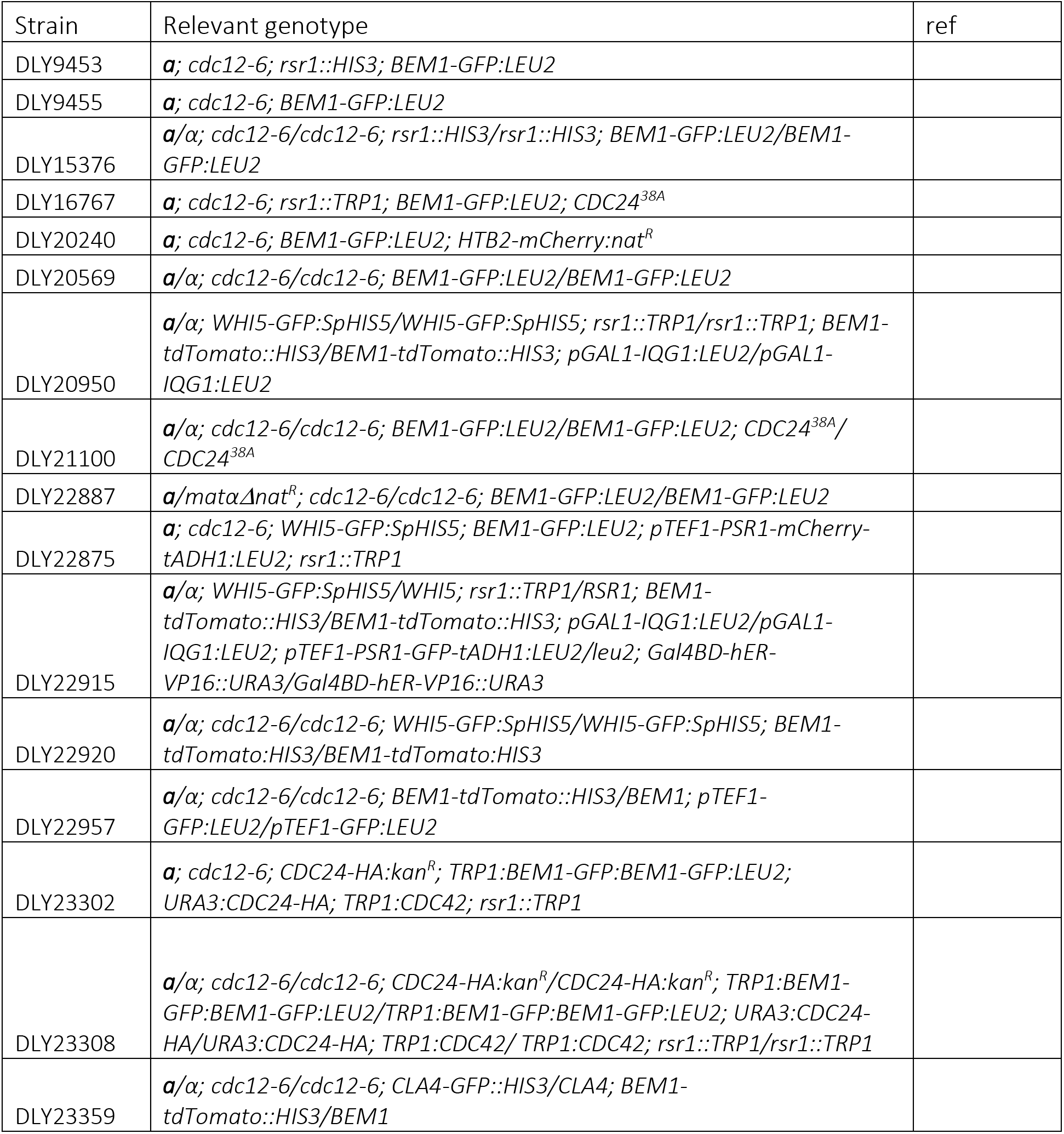

### Cell growth, hydroxyurea treatment, and timelapse imaging conditions

Cells were grown in liquid complete synthetic media (CSM, MP Biomedicals) with 2% dextrose at 24°C overnight until they reached log phase (5×10^6^ cells/mL). cdc12-6 cultures were shifted to 37°C and treated with 200 mM hydroxyurea (Sigma) for 1 h to protect cells from subsequent phototoxicity during imaging (Howell et al., 2012). Cells were pelleted, washed with and released into fresh media at 37°C for an additional 1 h (for imaging of the second cell cycle) or 3 h (for imaging of the third cell cycle). Cells were then harvested by centrifugation and mounted on a 37°C slab composed of CSM solidified with 2% agarose (Denville Scientific, Inc.) prior to imaging.

Images in Figure 3A were taken with live cell imaging, the cells were imaged at 37°C on Axio Observer.Z1 (Zeiss) with Pecon XL S1 incubator and control modules, a X-CITE 120XL metal halide fluorescence light source, and a 100x/1.46 (Oil) Plan Apochromat objective controlled by MetaMorph 7.8 (Universal Imaging). Images were captured with a Photometrics Evolve back-thinned EM-CCD camera. The fluorescence light source was set to 50% of the maximal output with a 2% ND filter. An EM-Gain of 750 and 200 ms exposure was set for the red channel (HTB2-mCherry), and an EM-Gain of 100 and 20 ms exposure was set for the Differential interference contrast (DIC) channel.

Images other than Fig. 3A were taken with confocal imaging, images were acquired with an Andor XD revolution spinning disk confocal microscope (Olympus) with a Yokogawa CsuX-1 5000 rpm disk unit and a 100x/1.4 U PlanSApo oil-immersion objective controlled by MetaMorph 7.8. 20 Z-stacks of 0.5 μm z-step were captured at 45-s intervals with Andor Ixon3 897 512 EMCCD camera (Andor Technology). The fluorescence light source was set to 6% of the maximal laser power for the 488 nm channel and 8% for the 561 nm channel. An EM-Gain of 200 and exposures of 250 ms were used.

Fluorescent images were deconvolved with SVI Huygens Deconvolution (Scientific Volume Imaging) and analyzed using Fiji (Schindelin et al., 2012). For deconvolution, a signal to noise ratio of 10 was used for live cell images and a ratio of 3 was used for Confocal images. Only cells that are not connected to neighboring cells were used for quantification to avoid cell pairs that might be connected from the previous cell cycle.

### Cell Fixation and membrane staining

To score the number of buds at the second cell cycle in Fig. 3H, cells were grown overnight at 24°C and log phase cultures (10^7^ cells/mL) were shifted to 37°C for 4 h. 1 mL cell culture was then harvested, spun down, and resuspended in 100 μL ice cold 10 μM FM4-64fx in water (Thermofisher Scientific) on ice. After 1 min staining, 1 mL ice cold 4% paraformaldehyde was added and the mixture was incubated on ice for 10 min. The cells were then washed twice with phosphate-buffered saline (PBS) and stored at 4°C. Images were then taken, and only cells that were budding from two connected compartments were counted.

### Fluorescence recovery after photobleaching (FRAP)

FRAP experiments were conducted on a DeltaVision Elite Deconvolution Microscope (Applied Precision) with a 100x/1.40 oil UPLSAPO100X0 1-U2B836 WD objective controlled by SoftWoRx 6.1 (Softworx Inc.). Images were captured with a Coolsnap HQ2 high resolution CCD camera. Photobleaching experiments were conducted on small-budded cycle 2 *cdc12-6* cells over-expressing cytoplasmic GFP. The bleaching 488 nm laser was used for 5 ms at 20% of maximal intensity. Cells were imaged with 50 ms exposure time, and 2×2 binning for three images before bleach and 15 images after bleach. Imaging interval was set automatically assuming 1 s half-time.

Images were analyzed using Fiji and MATLAB (Mathworks). Fluorescence signal at the bleach site and at sites in the mother and daughter compartments equidistant to the bleach site were averaged within a 3 μm diameter and normalized with an unbleached cell and the background fluorescence nearby using the formula:

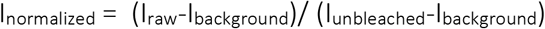

Normalized intensity at the mother site was fit to an exponential decay ae^−kt^+c, normalized intensity at the bleach site was fit to an exponential recovery -ae^−kt^+c, and normalized intensity at the daughter site was fit to a linear combination of the two ae^−kt^+be^ct^ + d. The recovery half-time can then be calculated by T_1/2_ = ln(2)/k.

### Simulated FRAP

The 3-dimensional geometry of a typical second-cycle *cdc12-6* cell was modeled by the closest point method described in (Ramirez et al., 2015). The cell shape was designated as the combination of a 6 μm diameter sphere and an ellipsoid with a long arm of 6 μm and a short arm of 2 μm, partially overlapped to create a neck of 2 μm diameter. The shape of the cell was modeled in Cartesian coordinates with the boundary of the cell interpolated with the closest grid points. The closest points were implemented with C++, and the main diffusion code was simulated by the implicit Euler method in MATLAB. The bleach was incorporated in the initial condition as a cylinder of of 1 μm diameter and zero intensity. “Fluorescence intensities” were the measured from the sum of z-stacks to mimic non-deconvolved microscopy images from the DeltaVision microscope.

### Polarity Models

We considered four polarity models in this study: a minimalistic mass-conserved activator-substrate (MCAS) model, a mechanistic model of the yeast polarity circuit, an extension of the MCAS model that incorporates an indirect substrate, and a minimalistic model that is no longer mass-conserved.

The minimalistic MCAS model considers the concentrations of two interconvertible forms of a protein (activator and substrate: u, v) in one spatial dimension (Fig. 1c)(Chiou et al., 2018). The protein can diffuse and convert between the two forms but is not synthesized or degraded:

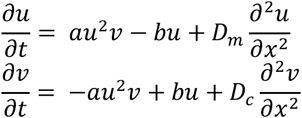

u enhances the conversion of v into more u through an implicit positive feedback loop modeled by the quadratic term au^2^v. u converts back to v in a first order process. u diffuses slowly relative to v. The parameters are:

| Description | Parameter | Value | reference |
| --- | --- | --- | --- |
| $u \rightarrow v$ | a | 1 | Chiou et al., 2018 |
| $v \rightarrow u$ | b | 1 | Chiou et al., 2018 |
| Diffusion constant of u | $D_m$ | 0.01 | Chiou et al., 2018 |
| Diffusion constant of v | $D_c$ | 1 | Chiou et al., 2018 |

The indirect-substrate model is similar to the minimalistic model except for the inclusion of a new species, the indirect-substrate v_i_ (Fig. 2c). u converts to v_i_ in a first-order process, and v_i_ converts to v in a first-order process. The differences from the minimalistic MCAS model are highlighted in bold:

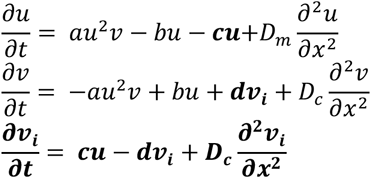

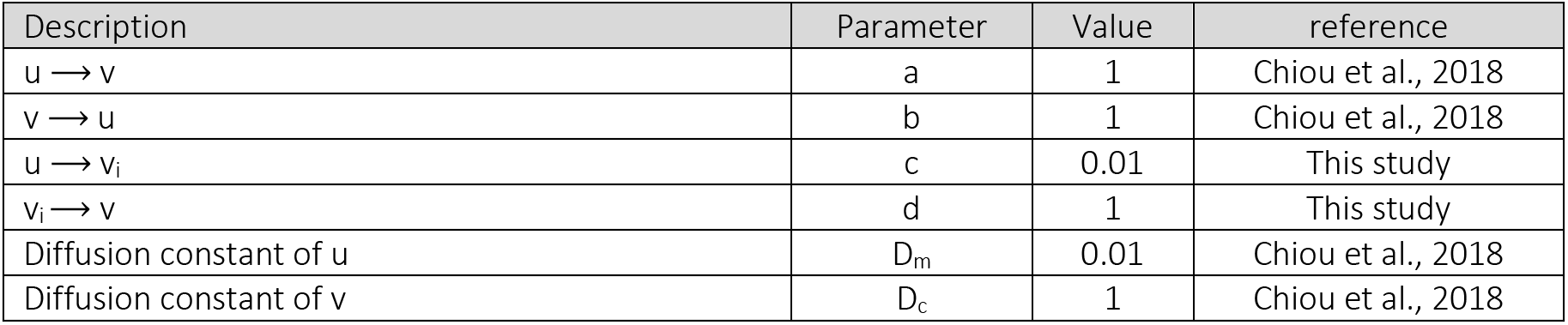

The mechanistic positive feedback model (Wu et al., 2015) is based on reactions assumed to occur with first order kinetics that account for various interconversions of Cdc42 and PAK-Bem1-GEF complexes. Cdc42 can interconvert between active GTP-bound (Cdc42T) and inactive GDP-bound (Cdc42D) states. Activation is catalyzed by GEF at the membrane, while inactivation is catalyzed by an implicit GAP. GDP-Cdc42 can also exchange between membrane (Cdc42D_m_) and cytoplasmic (Cdc42D_c_) forms (in cells this is catalyzed by GDP-dissociation Inhibitor or GDI). The PAK-Bem1-GEF complex (here called BemGEF) can similarly exchange between membrane (BemGEF_m_) and cytoplasmic (BemGEF_c_) forms, and in all cases membrane species diffuse mush less than cytoplasmic species. Positive feedback occurs due to reversible binding of BemGEF to Cdc42T, generating the complex BemGEF42 at the membrane. This leads to accumulation of GEF at sites with elevated GTP-Cdc42, which promotes local activation of more Cdc42. These reactions are modeled as:

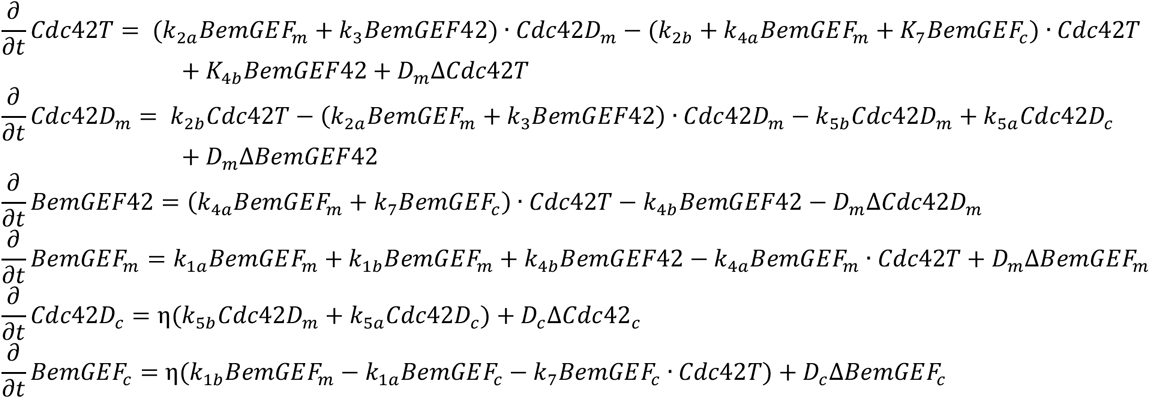

| Description | Parameter | Value | unit | reference |
| --- | --- | --- | --- | --- |
| $BemGEF_c \rightarrow BemGEF_m$ | $k_{1a}$ | 10 | $s^{-1}$ | Goryachev and Pokhilko, 2008 |
| $BemGEF_m \rightarrow BemGEF_c$ | $k_{1b}$ | 10 | $s^{-1}$ | Goryachev and Pokhilko, 2008 |
| $Cdc42D_m + BemGEF_m \rightarrow Cdc42T_m + BemGEF_m$ | $k_{2a}$ | 0.16 | $s^{-1}$ | Howell et al., 2009 |
| $Cdc42T \rightarrow Cdc42D_m$ | $k_{2b}$ | 1.75 | $\mu M^{-1} s^{-1}$ | Wu et al., 2015 |
| $Cdc42D_m + BemGEF42 \rightarrow Cdc42T + BemGEF42$ | $k_3$ | 0.35 | $s^{-1}$ | Howell et al., 2009 |
| $BemGEF + Cdc42 \rightarrow BemGEF42$ | $k_{4a}$ | 10 | $\mu M^{-1} s^{-1}$ | Goryachev and Pokhilko, 2008 |
| $BemGEF42 \rightarrow BemGEF + Cdc42T$ | $k_{4b}$ | 10 | $\mu M^{-1} s^{-1}$ | Goryachev and Pokhilko, 2008 |
| $Cdc42D_c \rightarrow Cdc42D_m$ | $k_{5a}$ | 36 | $s^{-1}$ | Kuo et al., 2014 |
| $Cdc42D_m \rightarrow Cdc42D_c$ | $k_{5b}$ | 0.65 | $s^{-1}$ | Kuo et al., 2014 |
| $BemGEF_c \rightarrow Cdc42T \rightarrow BemGEF42$ | $k_7$ | 10 | $\mu M^{-1} s^{-1}$ | Goryachev and Pokhilko, 2008 |
| Diffusion constant on the membrane | $D_m$ | 0.0025 | $\mu m^2 s^{-1}$ | Goryachev and Pokhilko, 2008 |
| Diffusion constant in the cytoplasm | $D_c$ | 10 | $\mu m^2 s^{-1}$ | Goryachev and Pokhilko, 2008 |
| Membrane to cytoplasm volume ratio | $\eta$ | 0.01 | | Goryachev and Pokhilko, 2008 |

In addition, the yeast polarity circuit contains a negative feedback loop due to multi-site phosphorylation of the GEF by the PAK, causing inactivation of the GEF (Kuo et al., 2014). Phosphorylation occurs when the PAK from one complex phosphorylates the GEF from another complex, which only happens when both complexes are bound to GTP-Cdc42. Dephosphorylation occurs only in the cytoplasm. The phosphorylated species, BemGEF*, can still exchange between cytoplasmic (BemGEF*c) and membrane (BemGEF*_m_) forms, and bind reversibly to Cdc42T (generating BemGEF*42). The addition of negative feedback leads to the differences highlighted in bold:

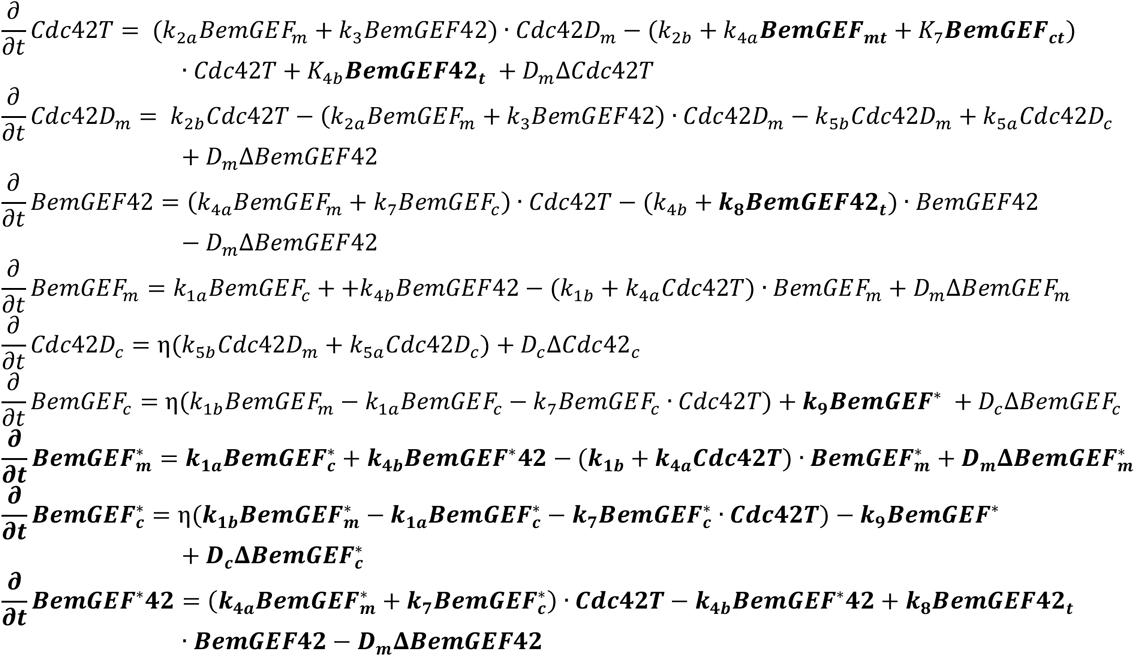

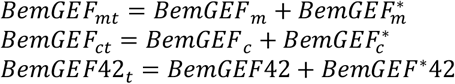

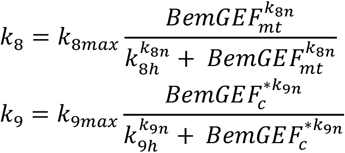

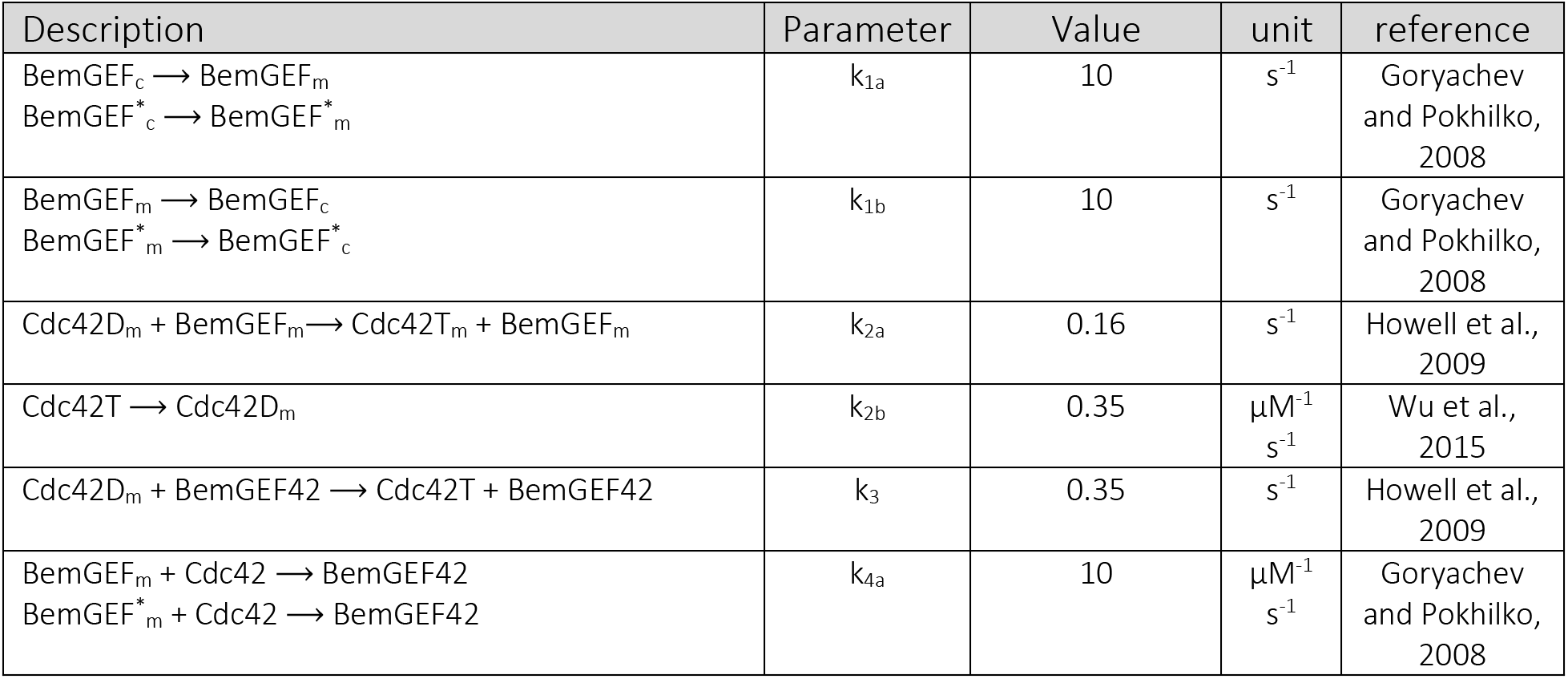
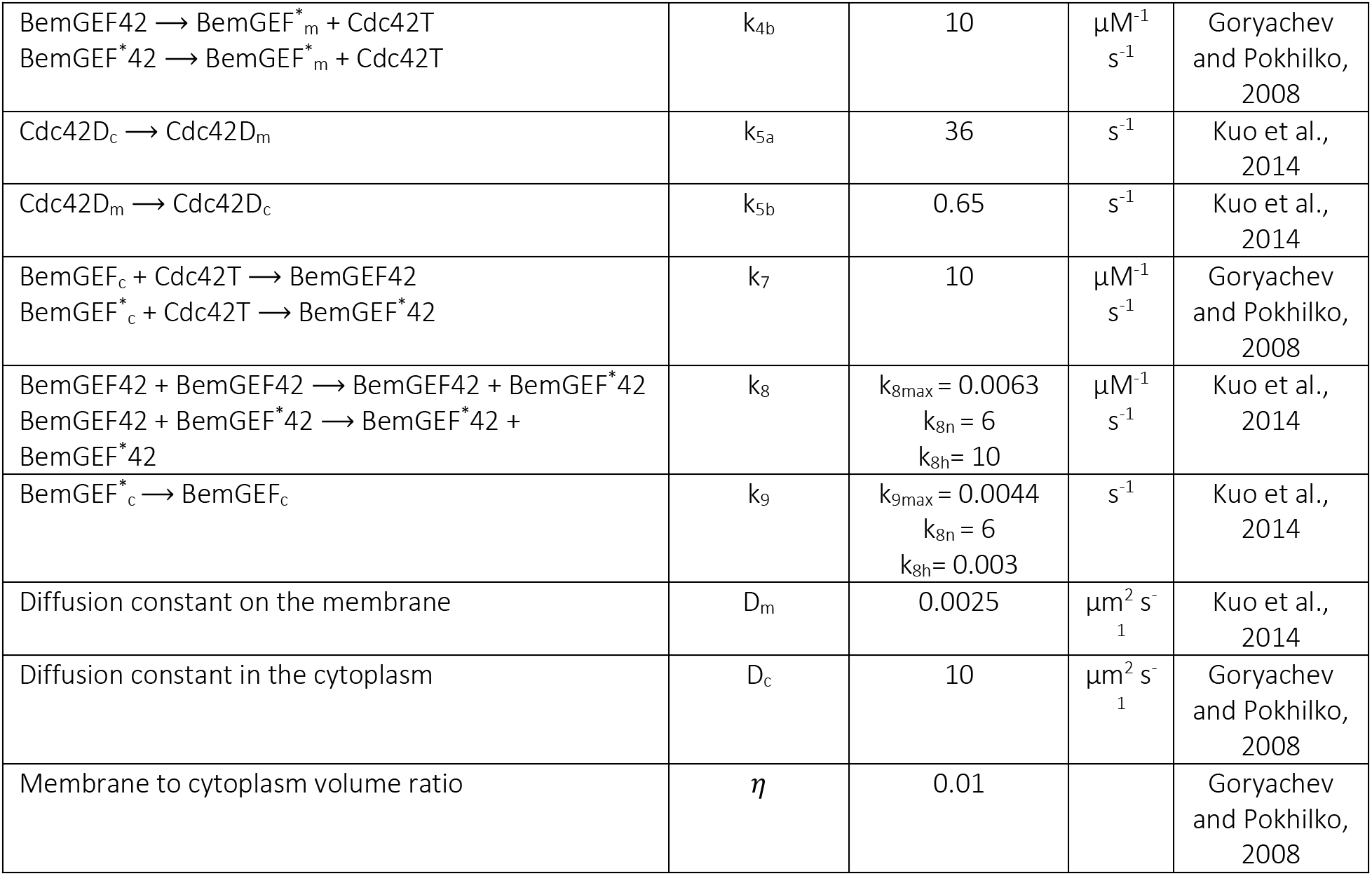

Note that the subscript “t” is used to denote the sum pf phosphorylated and unphosphorylated species, which can both undergo reversible binding to either membranes or GTP-Cdc42. Also, although only unphosphorylated species can act as GEFs, both phosphorylated and unphosphorylated complexes can act as PAKs to phosphorylate other GEFs. Multi-site phosphorylation and dephosphorylation are assumed to occur in an ultrasensitive manner.

Simulations of the MCAS models were done on MATLAB. Simulations of conceptual models were done on 1-dimensional domains with spatial resolution of 500 grid points. Finite differences were used with the linear diffusion being treated implicitly and the nonlinear reaction term explicitly in the time stepping. Mechanistic models were simulated on 2-dimensional domains with 200×200 grid points. All simulations proceeded with adaptive time stepping according to relative error in the reaction term. The MATLAB code used for simulations is provided in Source Code Files.

## Acknowledgements

We thank Tim Elston, Nick Buchler, Stefano Di Talia, Amy Gladfelter, Masayuki Onishi, and members of the Lew lab for comments on the manuscript. Thanks to Trevin Zyla for help with yeast strain construction. This work was funded by NIH/NIGMS grant R35GM122488 to D.J.L.

